# The Integrated Stress Response effector GADD34 is repurposed by neurons to promote stimulus-induced translation

**DOI:** 10.1101/2023.07.12.548606

**Authors:** Mauricio M. Oliveira, Muhaned Mohamed, Megan K. Elder, Keylin Banegas-Morales, Maggie Mamcarz, Emily H. Lu, Ela A.N. Golhan, Nishika Navrange, Snehajyoti Chatterjee, Ted Abel, Eric Klann

## Abstract

Neuronal protein synthesis is required for long-lasting plasticity and long-term memory consolidation. Dephosphorylation of eukaryotic initiation factor 2α is one of the key translational control events that is required to increase *de novo* protein synthesis that underlies long-lasting plasticity and memory consolidation. Here, we interrogate the molecular pathways of translational control that are triggered by stimulation of neurons with brain-derived neurotrophic factor (BDNF), which results in eIF2α dephosphorylation and *de novo* protein synthesis. Primary rodent neurons exposed to BDNF displayed elevated translation, but not transcription, of GADD34, which facilitates eIF2α dephosphorylation and subsequent *de novo* protein synthesis. Furthermore, GADD34 requires G-actin generated by cofilin to dephosphorylate eIF2α and enhance protein synthesis. Finally, GADD34 is required for the BDNF-induced translation of synaptic plasticity-related proteins. Overall, we provide evidence that neurons repurpose GADD34, an effector of the Integrated Stress Response, as an orchestrator of rapid increases in eIF2-dependent translation in response to plasticity-inducing stimuli.

## Introduction

In 1964, Flexner and colleagues^1^ demonstrated that the long-term memory formation relies on the synthesis of new proteins. Since then, increasing evidence has demonstrated that rapid increases in neuronal *de novo* protein synthesis is critical for memory consolidation^2^. Moreover, dysregulation of protein synthesis is a central feature in several neurodevelopmental and neurodegenerative disorders^3–13^. These observations have generated intense interest regarding how neurons control translation in response to activity, and how this contributes to brain function, including cognitive function.

*Cis*-regulatory elements present at untranslated regions (UTRs) flanking the main open reading frame (ORF) localize mRNAs throughout subcellular compartments, conferring the evolutionary advantage of rapid proteome modifications locally without the need for protein trafficking^14–20^. In neurons, this becomes even more pronounced, due to their extensive projections; 90% of the neuronal proteome is located at these projections^21^. In recent years, great effort has been directed toward identifying the mRNAs that are translated in dendrites and axons, which has revealed mRNAs that encode structural plasticity-related proteins located to synapses, such as cytoskeleton modifiers, post-synaptic density scaffolding components and ribosomal proteins^22–28^. Overall, these findings suggest that *de novo* protein synthesis coordinates a number of processes involved in synaptic development and transmission, such as ion channel insertion/depletion in the postsynaptic density^29–32^, dendritic spine expansion^33–36^, pre-synaptic arbor development^37–39^, axon growth^39–43^, and neurotransmitter release^44-46^.

The translation of both somatic and dendritic mRNAs is enhanced in response to extracellular stimuli, including neurotransmitters and neurotrophins. These include brain-derived neurotrophic factor (BDNF), which is released in a calcium-dependent manner^47, 48^, binds to postsynaptic tyrosine kinase receptors, and triggers at least two major pathways that can promote protein synthesis: extracellular-regulated kinase (ERK) and mechanistic target of rapamycin complex 1 (mTORC1)-mediated pathways^49–52^, that promote translation of mRNAs via eIF4E and eEF2^51, 53–55^. BDNF also can induce protein synthesis-dependent long-term potentiation in the hippocampus^49, 56–61^, dendritic spine expansion^35, 62^ and calcium-dependent release of neurotransmitters^44, 60^. Thus, release of BDNF can trigger synaptic signaling to increase *de novo* protein synthesis.

Despite the identification of mRNAs localized to dendrites, how these quiescent mRNAs become translated in response to plasticity-inducing stimuli is still an open question. Translation is a multi-step and -factorial process with an overwhelming number of regulatory checkpoints. Neuronal translation initiation is regulated at two main checkpoints: the phosphorylation of the eukaryotic initiation factor 2 on its alpha subunit (eIF2α-P) and the phosphorylation of eIF4E/4E-binding protein 2 (4E-BP2)^63^. eIF4E has been implicated in plasticity-related events, stimulating the translocation of ribosomes from the dendritic shaft to spines^64^ and promoting synthesis of proteins that are known to be important for synaptic and memory consolidation^65^. eIF4E mRNA and protein also were shown to be enriched in dendritic spines during memory formation^66, 67^.

The first step of translation initiation requires eIF2, which is responsible for carrying loaded ^Met^tRNA-Met to ribosomes. The transfer of the amino acid from tRNA to the nascent peptide chain requires guanosine triphosphate (GTP) to be bound by eIF2, which is promoted by the guanosine exchange factor eIF2B^68–72^. When phosphorylated on its α subunit (eIF2α-P), eIF2 blocks the exchange activity of eIF2B and inhibits global translation^73^. The phosphorylation of eIF2α is critical for the Integrated <u>S</u>tress <u>R</u>esponse (ISR), which is triggered by different cellular stressors. These stressors converge to rapidly increase eIF2α-P levels via activation of protein kinases that sense stress, resulting in an inhibition of general protein synthesis to conserve energy until the cellular stress is resolved. The reduction of protein synthesis coincides with the translation of a subset of mRNAs with upstream open reading frames (uORFs) in their 5’ untranslated region (UTR)^74^. One such mRNA, *PPP1R15A*, encodes GADD34, a scaffolding protein that acts in a negative feedback loop to the ISR. GADD34 is a protein phosphatase 1 (PP1)-targeting protein that directly binds to eIF2α-P, thereby promoting its dephosphorylation and restoration of protein synthesis after stress resolution^75^.

Decreased levels of eIF2α-P are associated with neuronal activity and memory formation. Long-term potentiation (LTP)^76^ and threat conditioning in the basolateral amygdala^77^ and hippocampus^78^ is associated with decreased levels of eIF2α-P. Moreover, it was shown that blocking the decrease in eIF2α-P following learning impairs long-term memory consolidation^79^. At the cellular level, eIF2α phosphorylation controls spine expansion^80^ and promotes the insertion of AMPAR in the post-synaptic density^29^. However, the mechanism responsible for the plasticity-induced downregulation of eIF2α-P remains unknown. We considered two main hypotheses: plasticity-inducing stimulation (1) decreases eIF2α kinase activity, resulting in reduced eIF2α-P and/or (2) increases eIF2α dephosphorylation via alteration of regulators of the phosphatase. Because the latter offers a direct and more rapid way of altering eIF2α-P (instead of kinase inhibition followed by passive dephosphorylation), we hypothesized that neuronal activation results in GADD34-mediated dephosphorylation of eIF2α, thereby increasing protein synthesis after neuronal activity. Herein, we provide evidence that neuronal stimulation with BDNF increases GADD34 expression in neurons, and that this increase relies on translational, rather than transcriptional, regulation. Moreover, BDNF increases the physical interaction between GADD34 and eIF2α in both soma and dendrites, and GADD34 expression converges with actin cytoskeleton dynamics that is required for the BDNF-induced increase in protein synthesis. Finally, GADD34 orchestrates the translation of numerous mRNAs related to neurotransmission-related metabolic functions as well as synaptic organization and activity. Overall, our findings support the notion that activity-dependent increases in neuronal protein synthesis rely on translation-dependent increases in GADD34 expression, revealing a novel molecular mechanism by which neuronal activation promotes rapid *de novo* translation.

## Results

### GADD34 is constitutively expressed in neurons

GADD34 has classically been described as a negative regulator of the ISR, expressed solely under stress conditions^75^. However, examination of a brain cell type-specific RNA-seq database^81^ showed that GADD34 mRNA is present in neurons, microglia, and astrocytes in the mouse brain (data not shown). These findings prompted us to determine whether GADD34 is constitutively expressed in neurons.

We first employed fluorescent *in situ* hybridization (FISH) RNA scope to verify presence and localization of *PPP1R15A* mRNA (which encodes GADD34) in primary neurons. We found that *PPP1R15A* mRNA is localized in both the soma (∼75% of total FISH signal) and dendrites (∼25%) of mature (*DIV* 14) mouse cortical neurons (Fig 1A-B). Treatment with the stressor thapsigargin showed that the signal was specific to *PPP1R15A* mRNA (Fig S1A-D). To study protein expression in the cell reliably (i.e. confirmation of specific antibody staining), we targeted GADD34 expression in a cell type-specific way, engineering a new viral tool that allowed the expression of an shRNA targeting *PPP1R15A* mRNA in a Cre-dependent manner (AAV9.EF1.mCherry.SICO-shRNA.PPP1R15A, Fig 1C). With this virus, we selectively decreased the expression of GADD34 in excitatory neurons employing a viral cocktail composed of AAV9.CamKIIα-Cre driver and AAV9.EF1.mCherry.SICO-shRNA.PPP1R15A (Fig 1D). Using this approach, we were able to transduce 31% of total neurons with the shRNA (Suppl Fig 1E), from which 78.6% express the shRNA (Suppl Fig 1F). We then evaluated whether GADD34 protein could be detected in neurons with immunocytochemistry. We found that GADD34 is expressed in both soma and dendrites of neurons, and that this expression was downregulated when neurons expressed the shRNA targeting GADD34 (Fig 1E-F). We further confirmed these results with western blot analysis of neurons transduced with the viral cocktail and found GADD34 protein expression was significantly reduced by the shRNA (Fig 1G). Surprisingly, we did not see significant alterations in eIF2α-P levels, suggesting that GADD34 plays minimal role in homeostatic maintenance of neuronal initiation (Fig 1H). Overall, these results confirmed that GADD34 is constitutively expressed in neurons.

**Figure 1.**
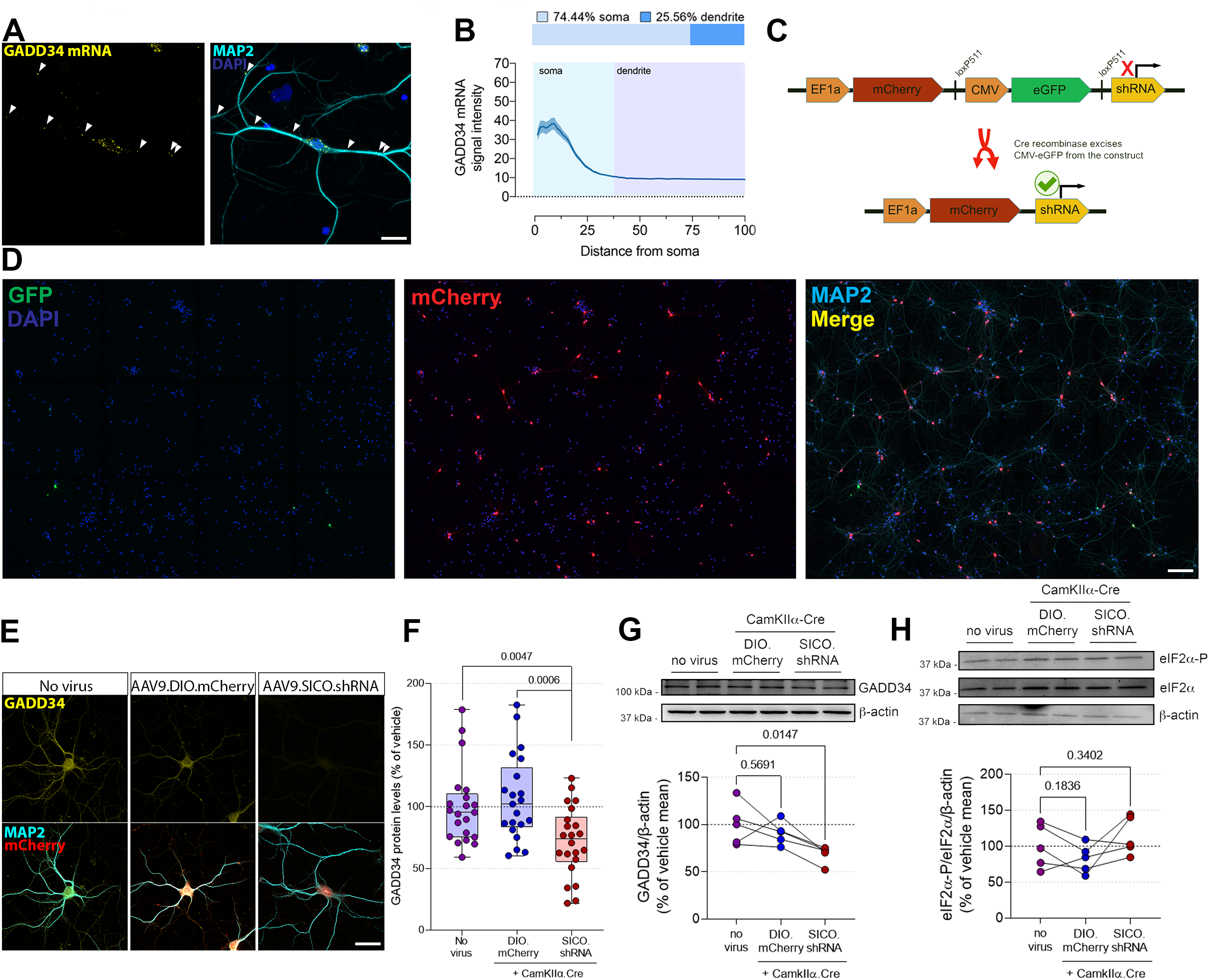
GADD34 is constitutively expressed in neurons. (**A**) Fluorescent *in situ* hybridization (FISH) targeting GADD34 mRNA (punctate signal). Green=FISH; Blue=DAPI; Cyan=MAP2. Scale bar=50 μm. (**B**) Upper panel: percentage of total mRNA localized to somatic versus dendritic compartments. Bottom panel: radial quantification of GADD34 mRNA puncta in neurons, from soma (light blue background) to dendritic portions (dark blue background) (n=3 independent cultures). (**C**) SICO.shRNA system for knockdown of GADD34. (**D**) Neurons transduced with AAV9.EF1α.mCherry.SICO-shRNA.PPP1R15A. Scale bar=200 μm. (**E**) Representative images of GADD34 immunostaining in neurons transduced with AAV9.EF1α.mCherry.SICO-shRNA.PPP1R15A. Yellow=GADD34; Cyan=MAP2; Red=mCherry. (**F**) Quantification of “E” (n=20-23 neurons from 2 independent cultures). One-way ANOVA followed by Dunnett *post-hoc* test. Scale bar=50 μm. (**G**) Western blot of cell lysates obtained from naïve neurons or co-transduced with AAV9.CamKIIα.Cre + AAV9.hSyn2.DIO.mCherry or AAV9.CamKIIα.Cre + AAV9.EF1α.mCherry.SICO-shRNA.PPP1R15A (n=5 independent cultures). Upper row=GADD34; lower row=β-actin. One-way ANOVA followed by Dunnet *post-hoc* test. (**H**) Western blot of cell lysates obtained from naïve neurons or co-transduced with AAV9.CamKIIα.Cre + AAV9.hSyn2.DIO.mCherry or AAV9.CamKIIα.Cre + AAV9.EF1α.mCherry.SICO-shRNA.PPP1R15A (n=5 independent cultures). Upper row=eIF2α-P; Middle row=total eIF2α; Lower row=β-actin. One-way ANOVA followed by Dunnet *post-hoc* test.

### BDNF increases GADD34 translation in neurons

GADD34 was previously shown to have its expression in neurons tied to plasticity-related transcription factors^82, 83^. Therefore, we hypothesized neuronal stimulation could increase GADD34 levels in primary neurons. To address this question, we induced translation-dependent activity in primary neurons using BDNF. We exposed primary neurons to BDNF (50 ng/ml) for 0, 5, 15, 30, 60 or 120 min and then probed for GADD34 protein levels via western blot. We found that GADD34 was significantly increased 60 min after exposure to BDNF and this increase persisted until at least 120 min, which was accompanied by a concomitant decrease in eIF2α-P levels (Fig 2A-C), suggesting that GADD34 could be a direct mediator of BDNF-induced decreases in eIF2α-P. These findings indicated that neuronal GADD34 expression can be induced by BDNF.

**Figure 2.**
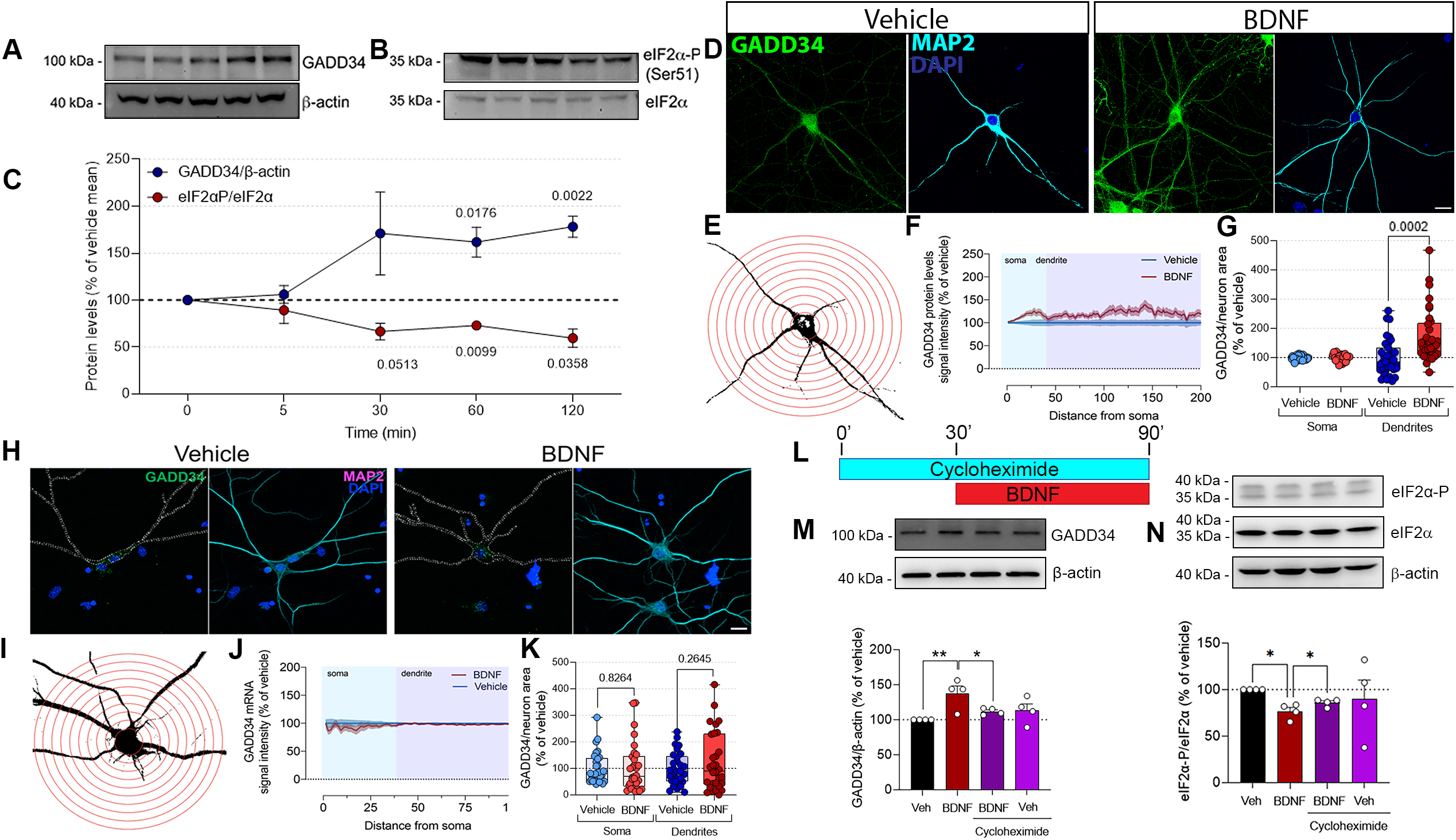
BDNF promotes transcription-independent increase in GADD34 protein levels. (**A**) Western blot probed for GADD34 (upper blot) or β-actin control (lower lane) using samples obtained from neurons exposed to BDNF at time points indicated. (**B**) Western blot probed for eIF2α-P (Ser51, upper lane) or total eIF2α (lower lane) using the same samples described in panel **A**. (**C**) Quantification of “A” and “B” (n=5 independent primary cultures). One-way ANOVA repeated measures followed by Dunnett *post-hoc* test. (**D**) Immunofluorescence staining for GADD34 in neurons. Green=GADD34; Cyan=MAP2; Blue=DAPI. Scale bar=50 μm. (**E**) Representation of the radial quantification of GADD34 expression in neurons. (**F**) Radial quantification of GADD34 expression in soma (light blue background) and in dendrites (dark blue background) of neurons exposed to either vehicle (blue line) or BDNF for 1 hr (red line). Light shade surrounding the lines represent SEM. (**G**) Total amount of GADD34 in either soma or dendrites of neurons exposed to either vehicle or 50 ng/ml BDNF for 1 hr. Statistical analysis was performed to compare differences in each compartment, independently (n=37-38 neurons/condition from 3 independent cultures). Unpaired *t* test. (**H**) RNA scope-based FISH staining against GADD34 mRNA in neurons. Green=FISH; Cyan=MAP2; Blue=DAPI. Scale bar=50 μm. (**I**) Representation of the radial quantification of *PPP1R15A* mRNA expression in primary neurons. (**J**) Radial quantification of *PPP1R15A* mRNA expression in soma (light blue background) and in dendrites (dark blue background) of neurons exposed to either vehicle (blue line) or BDNF (red line) for 1 hr. Light shade surrounding the lines represent SEM. (**K**) Total amounts of *PPP1R15A* mRNA in either soma or dendrites of neurons exposed to either vehicle or BDNF for 1 hr (n=33-34 neurons/condition from 3 independent cultures). (**L**) Representation of timeline for experiment with cycloheximide and BDNF. (**M**) Western blot using samples obtained from neurons treated as described in panel “L” (n=4 independent cultures). Upper row=GADD34; lower row=β-actin. Two-Way ANOVA followed by Tukey’s *post-hoc* test, *=p<0.05; **=p<0.01. Mean ± SEM. (**N**) Western blot using samples obtained from neurons treated as described in panel “L” (n=4 independent cultures). Upper row=eIF2α-P; middle row=total eIF2α; lower row=β-actin. Two-Way ANOVA followed by Tukey’s *post-hoc* test, *=p<0.05. Mean ± SEM.

Our results and previous data^84^ showed that GADD34 mRNA (*PPP1R15A*) is present both in soma and dendrites of neurons (Fig 1). We then examined whether the increase in GADD34 levels was compartmentalized. We first determined total levels of GADD34 in somatic versus dendritic compartments of neurons treated with BDNF (Fig 2D) and performed a radial quantification of GADD34 staining in dendritic arbors (Fig 2E). Our results indicated that GADD34 protein levels are increased in dendrites, but not the soma, 1 hr after stimulation with BDNF (Fig 2F-G). We then asked whether the increase in GADD34 expression was due to transcriptional modulation, as BDNF is known to increase ATF4 in neurons^85^. We observed no differences in *PPP1R15A* mRNA levels after exposure to BDNF (Fig 2H-K). Moreover, when we pre-treated neurons with cycloheximide (Fig 2L), a potent translation inhibitor, BDNF failed to increase GADD34 levels (Fig 2M), as well as to decrease eIF2α-P (Fig 2N). Thus, these data indicate that the increase in BDNF-induced increase in GADD34 levels relies on translational, rather than transcriptional regulatory mechanisms.

Previous ribosome foot printing data indicate that *PPP1R15A* mRNA is bound to ribosomes, which are preferentially stalled at the uORF2 in the 5’UTR of the mRNA (Fig 3B)^84^. We combined this observation with our data (Fig 2M) and hypothesized that GADD34 protein levels are increased by BDNF via regulation at translational level. To test this hypothesis, we performed translating ribosome affinity purification (TRAP) to investigate changes in the amount of *PPP1R15A* mRNA loaded onto ribosomes of excitatory neurons exposed to BDNF (Fig 3A)^86^. We transduced primary neurons with a viral cocktail containing AAV9.CamKIIα-Cre and AAV9.CAG.FLEX.L10a-EGFP, to drive L10a-EGFP expression in excitatory neurons, which resulted in transduction of ∼75% of neurons (Fig S2A-B). Real time (RT)-PCR demonstrated that the amount of *PPP1R15A* bound to ribosomes was significantly increased in response to BDNF (Fig 3C, red points). Curiously, when we examined total *PPP1R15A* mRNA, we found a (non-significant) trend for enhanced levels following BDNF, also suggesting a transcriptional event in response to BDNF (Fig. 3C, blue points). However, it should be noted that this increase could also arise from either inhibitory neurons or glial cells that are present in the primary cultures we utilized.

**Figure 3.**
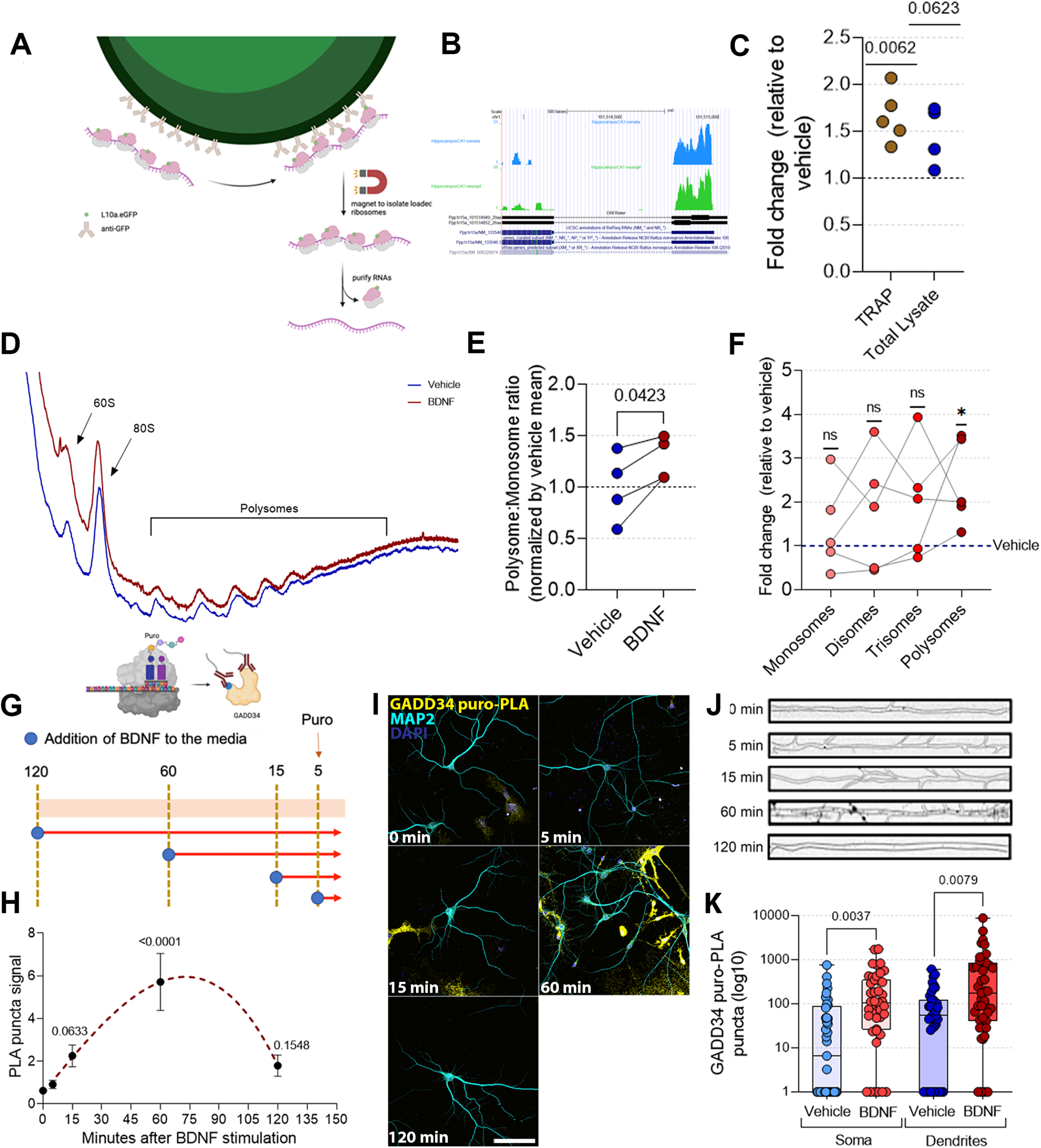
BDNF increases GADD34 translation in neurons. (**A**) Schematics of the workflow for TRAP-qPCR. (**B**) Ribo-seq data from Glock et al indicating stalled ribosomes at the uORFs of *PPP1R15A* mRNA in neuronal soma and dendrites. (**C**) TRAP-qPCR and qPCR (total lysate) analysis of *PPP1R15A* normalized by GAPDH (n=4-5 independent cultures). Wilcoxon rank test. (**D**) Representative polysome fractionation from neurons exposed to either vehicle (red) or BDNF for 1h (blue). (**E**) Polysome/monosome ratio. Paired *t* test, * = p<0.05. (**F**) Quantification of loaded *PPP1R15A* mRNA in different fractions of polysome profiling, assessed by RT-PCR (n=5 independent primary cultures). Blue dashed line represents vehicle, normalized to 1 in every fraction. Statistical analyses were performed comparing the mean variations per fraction. Wilcoxon rank test, ns=non-significant, *=p<0.05. (**G**) Puro-PLA experimental design. (**H**) Quantification of total puro-PLA signal in neurons exposed to BDNF in time points indicated (n=35-36 neurons from 3 independent cultures). One-way ANOVA repeated measures followed by Dunnett *post-hoc* test. (**I**) Representative images of puro-PLA from the experiment described in panel **H** Green=Puro-PLA; Red=MAP2; Blue=DAPI. Scale bar=150 μm. (**J**) Representative images of dendritic signal from the puro-PLA time course experiment. PLA signal is represented as black dots. (**K**) Quantification of GADD34 puro-PLA signal at the 60 min time point of exposure to BDNF, isolating somatic and dendritic compartments (n=35-36 neurons from 3 independent cultures). Unpaired *t* test.

To gain deeper knowledge on *PP1R15A* translation regulation, we performed a polysome fractionation with sucrose gradients, collected fractions representing monosomes, disomes, trisomes or heavy polysomes (4 or more), and measured total *PPP1R15A* mRNA per fraction. Our data indicated that the occupancy of *PPP1R15A* in the heavier fractions was higher when cells were exposed to BDNF, suggesting higher translation rates (Fig. 3D-F). Finally, we used proximity ligand assay (PLA) coupled to puromycilation (puro-PLA)^87^ to visualize amounts and localization of newly synthesized GADD34 in neurons (Fig 3G; Fig S2C-D). To determine whether *PPP1R15A* mRNA is more translated after BDNF exposure, we treated neurons for 0, 5, 30, 60 and 120 min with BDNF, and added puromycin during the last 5 minutes, to obtain a snapshot of GADD34 translation (Fig 3G). Strikingly, the synthesis of GADD34 peaked 60 minutes after exposure to BDNF (Fig 3H). Furthermore, compartment-specific analysis showed elevated GADD34 translation after exposure to BDNF in both soma and dendrites (Fig 3I-K). Altogether, these results demonstrate that the increase in GADD34 levels induced by BDNF are due to translational, rather than transcriptional, regulation.

### BDNF promotes GADD34-eIF2α interactions and GADD34-dependent protein synthesis

Previous work demonstrated that neuronal activity increases protein synthesis via reduced eIF2α-P^76, 77^. Therefore, we asked whether GADD34 was required for the BDNF-induced decrease in eIF2α-P and increase in protein synthesis. We first determined whether BDNF could promote interactions between GADD34 and eIF2α. To assess this, we used PLA, this time targeting GADD34 and eIF2α. We found that the PLA was a reliable approach to measure GADD34-eIF2α interactions, as thapsigargin significantly increased the PLA signal (Fig S3A-B). To obtain a spatiotemporal analysis of GADD34-eIF2α interactions, we exposed neurons to BDNF for 0, 5, 15, 30 and 60 min prior to fixation and subsequent PLA analysis (Fig. 4A-C). We found that the GADD34-eIF2α interactions peak at 60 min (Fig 4C, black dashed line). Notably, we observed that the PLA signal peaked at 30 min in the dendritic compartment (Fig 4C, red dashed line), whereas in the soma the peak was at later time points (Fig 4C, blue dashed line) after BDNF stimulation. Compartment-specific analysis of the PLA signal induced by BDNF indicated that the GADD34-eIF2α interaction was increased after 60 min BDNF stimulation in both soma and dendrites (Fig 4D-E). These findings indicate that BDNF increases GADD34-eIF2α interactions.

**Figure 4.**
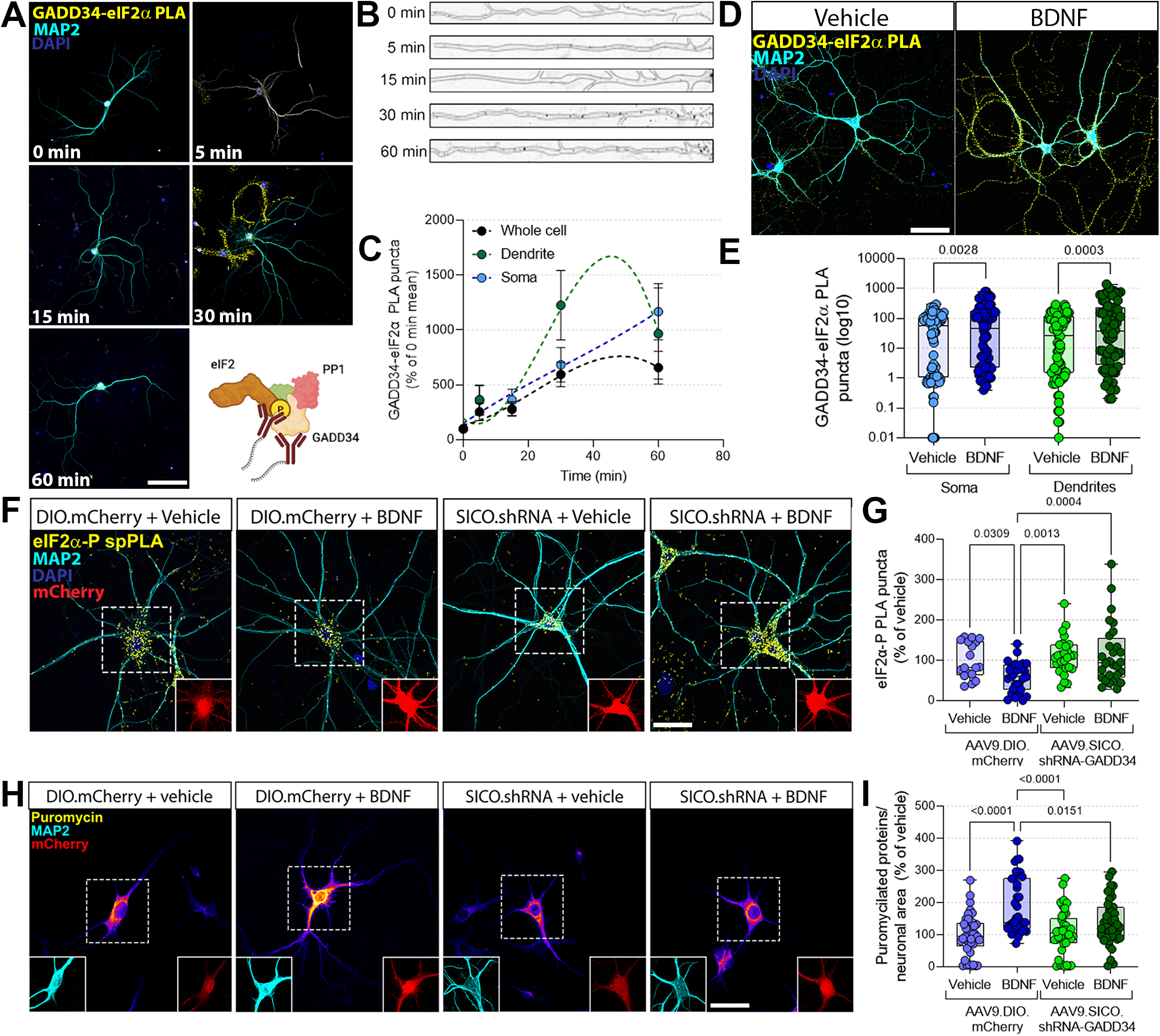
BDNF promotes GADD34-eIF2α interactions and GADD34-dependent protein synthesis. (**A**) Representative images of the PLA for GADD34-eIF2α in neurons exposed to BDNF in timepoints indicated. Green=PLA; Red=MAP2; Blue=DAPI. Insets are representative of the somatic PLA signal, represented by the black dots. Scale bar=100 μm. (**B**) Representative image of dendritic GADD34/eIF2α PLA signal over the time course of BDNF. (**C**) Quantification of “A” (N=22-24 neurons, from 2 independent cultures). Dashed red line with black dots represents total PLA signal normalized by neuronal area. Blue line and dots represent somatic signal. Continuous red line and dots represent dendritic PLA signal. One-way ANOVA repeated measures followed by Dunnett’s *post-hoc* test. (**D**) Representative images of GADD34-eIF2α PLA in neurons. Green=PLA; Red=MAP2; Blue=DAPI. Scale bar=100 μm. (**E**) Quantification of “D” (N=50 neurons from 3 independent cultures). Statistical analysis was performed comparing each compartment independently. Unpaired *t* test. (**F**) Representative images of eIF2α-P levels measured using spPLA in neurons transduced with either AAV9.EF1α.mCherry.SICO-shRNA.PPP1R15A or control AAV9.hSyn.DIO.mCherry. Cyan=MAP2; Red=mCherry; Yellow=eIF2α-P. Scale bar=50 μm. (**G**) Quantification of “F” (n=18-29 neurons/condition from 3 independent cultures). Two-Way ANOVA followed by Tukey’s *post-hoc* test. (**H**) Representative images of *de novo* protein synthesis measured using *in situ* SUnSET in neurons transduced with either AAV9.EF1α.mCherry.SICO-shRNA.PPP1R15A or control AAV9.hSyn.DIO.mCherry. Cyan=MAP2; Red=mCherry; Pseudo color=puromycin. Scale bar=50 μm. (**I**) Quantification of “F” (n=28-41 neurons/condition from 3 independent cultures). Two-Way ANOVA followed by Tukey’s *post-hoc* test.

To determine whether GADD34 is also required for BDNF-dependent increases in neuronal protein synthesis, we transduced neurons with a viral cocktail containing AAV9.CamKIIα-Cre and either AAV9.EF1.mCherry.SICO-shRNA.PPP1R15A or AAV9.hSyn.DIO.mCherry (control). We then tested whether GADD34 mediated the BDNF-dependent decrease in eIF2α-P. We found that decreasing GADD34 expression blocked eIF2α dephosphorylation (Fig. 4F-G) as well as the increase in protein synthesis by BDNF (Fig 4H-I) as assessed by puromycilation^88^. (Fig S3C). Collectively, our data indicate that GADD34 is necessary for the BDNF-induced increase in protein synthesis via eIF2α dephosphorylation.

### GADD34 mediates the BDNF-induced translation of synaptic plasticity-related mRNAs

Thus far, our data suggest that GADD34 is required for BDNF-induced protein synthesis. Therefore, we proceeded to determine the identity of the mRNAs whose translation is regulated in a GADD34-dependent manner with TRAP-sequencing, which allowed us to identify mRNAs specifically in excitatory neurons. We transduced neurons harboring *flox* inserts in the first two exons of GADD34 with AAV9.CamKIIα.Cre (henceforth called “KO” cells) and AAV9.CamKIIα.eGFP-L10a. As a control for the Cre recombinase (henceforth termed “WT” cells), we used AAV9.CamKIIα.mCherry (**Suppl. Fig. 4A**). At *DIV* 14, these neurons were exposed to BDNF for 1h and samples collected, and ribosomes purified using GFP antibodies (see Methods). Using this method, we found that 784 mRNAs were differentially expressed (DEGs) in the total lysate of WT cells (425 upregulated, 359 downregulated by BDNF) (Fig. 5A), whereas 3815 were DEGs in the TRAP-purified samples (1878 upregulated, 1937 downregulated by BDNF) (Fig. 5B). Furthermore, we found no significant outliers among samples using both principal component analysis (PCA; Suppl. Fig. 4B) and unsupervised clustering (Suppl. Fig. 4C). When we measured the correlation between total mRNA (i.e. transcriptional + translational changes) log_2_FC versus TRAP log2FC (i.e. translational changes), we found a small correlation (Pearson’s correlation coefficient = 0.1481) (Fig 5C), indicating that changes in the pool of mRNAs being differentially translated after 1h of BDNF exposure were not due to transcriptional changes. Of note, the analysis of DEGs obtained showed an enrichment for markers of excitatory neurons (*Camk2a, Grin1, Slc17a5* and *Slc17a7*) in the purified samples over total lysate, as well as a depletion of markers of inhibitory neurons (*Pvalb, Gad1*) and glial cells (*Gfap, Cnp*) (Suppl. Fig. 4D).

**Figure 5.**
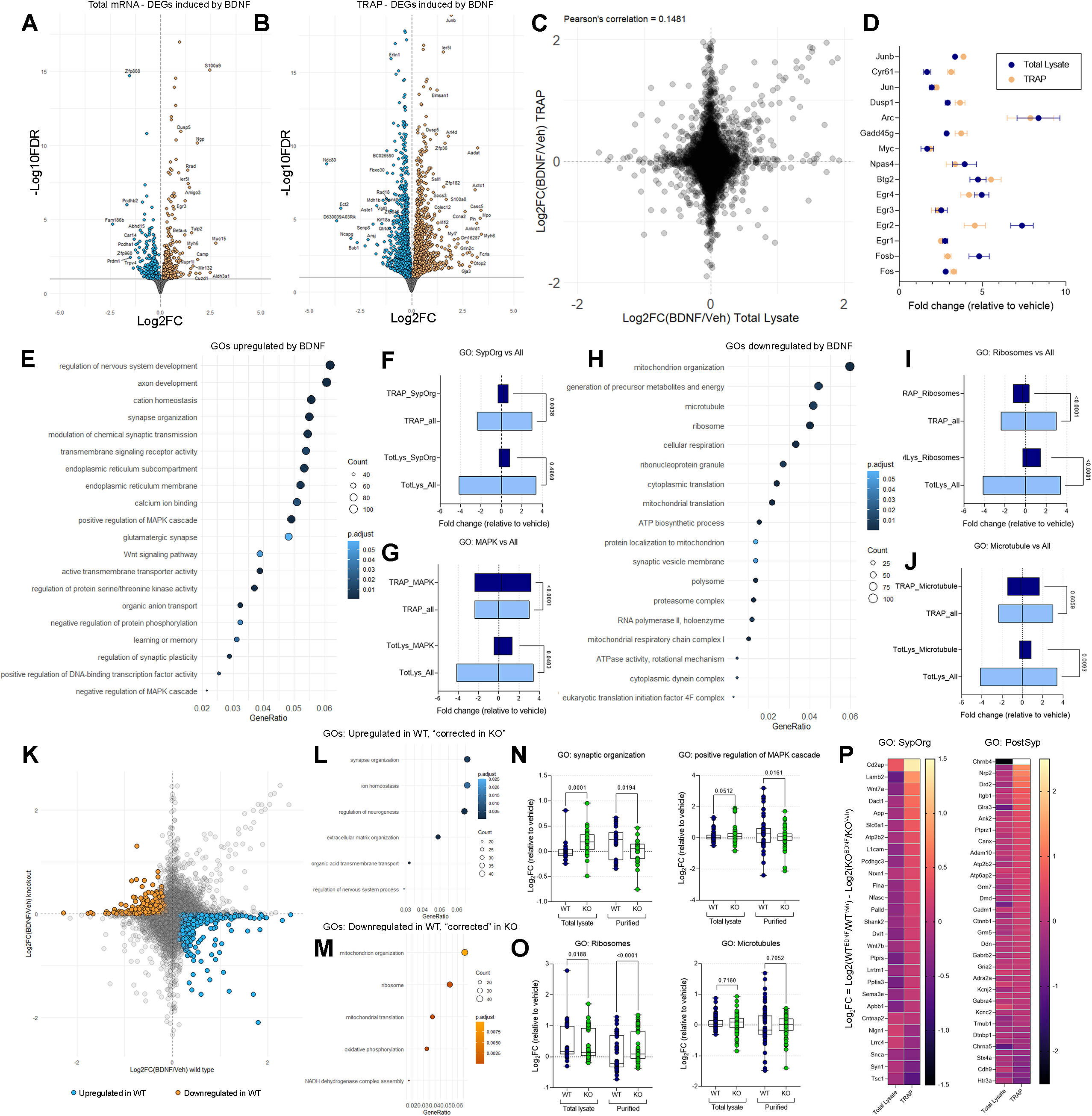
GADD34 mediates the BDNF-induced translation of mRNAs that encode synaptic plasticity-related proteins. (**A**) Volcano plot demonstrating log_2_FC of upregulated (orange) and downregulated (blue) mRNAs in total mRNA fraction. (**B**) Volcano plot demonstrating log_2_FC of upregulated (orange) and downregulated (blue) mRNAs in the TRAP purified fraction. (**C**) Correlation plot between Log_2_FC of total mRNA fraction (x-axis) versus TRAP fraction (y-axis). Each black dot is an individual mRNA. (**D**) Fold changes of IEGs induced by BDNF both in total fraction (blue) and TRAP fraction (beige). (**E**) GO analysis of DEGs significantly upregulated by BDNF. Size of circle indicates counts of mRNAs in each group, color indicates adjusted *p* value of that GO group. (**F**) Comparison of log_2_FC of mRNAs belonging to the “synapse organization” GO and all mRNAs identified in total mRNA or TRAP fractions. Gene ratio = number of genes identified in each group divided by the amount of background genes. Unpaired *t* test. (**G**) Comparison of log_2_FC of mRNAs belonging to the “positive regulation of MAPK cascade” GO and all mRNAs identified in total mRNA or TRAP fractions. Gene ratio is the same as in **F**. Unpaired *t* test. (**H**) GO analysis of DEGs significantly downregulated by BDNF. Size of circle indicates counts of mRNAs in each group, color indicates adjusted *p* value of that GO group. (**I**) Comparison of log_2_FC of mRNAs belonging to the “ribosome” GO and all mRNAs identified in total mRNA or TRAP fractions. Gene ratio is the same as in **F.** Unpaired *t* test. (**J**) Comparison of log_2_FC of mRNAs belonging to the “microtubule” GO and all mRNAs identified in total mRNA or TRAP fractions. Gene ratio is the same as in **F**. Unpaired *t* test. (**K**) Correlation plot between Log_2_FC of wild type (x-axis) versus KO cells (y-axis). Each dot is an individual mRNA. Orange dots indicate mRNAs that are downregulated in WT, corrected in KO. Blue dots indicate mRNAs that are upregulated in WT, corrected in KO. (**L**) GO analysis of DEGs significantly upregulated by BDNF, corrected in KO. Size of circle indicates counts of mRNAs in each group, color indicates adjusted *p* value of that GO group. (**M**) GO analysis of DEGs significantly downregulated by BDNF, corrected in KO. Size of circle indicates the total count of mRNAs in each group, color indicates adjusted *p* value of that GO group. (**N**) Comparison of log_2_FC WT vs log_2_FC KO of mRNAs belonging to the “synaptic organization” or “positive regulation of MAPK cascade” GOs. Comparisons were made in total and TRAP fractions. Unpaired *t* test. (**O**) Comparison of log_2_FC WT vs log_2_FC KO of mRNAs belonging to the “ribosome” or “microtubule” GOs. Comparisons were made in total and TRAP fractions. Unpaired *t* test. (**P**) Heatmaps comparing the log_2_FC of mRNAs belonging to either “synaptic organization” or “post synapse”. Comparisons were performed in total and TRAP fractions.

We first determined the expression of immediate early genes (IEGs) induced by BDNF in WT cells and found robust increases in all IEGs investigated in both TRAP and total mRNA fractions, showing expected plasticity-related transcriptional activation (Fig. 5D). Gene ontology (GO) analysis of the DEGs upregulated by BDNF revealed an enrichment of GOs related to neuronal development, synaptic organization and signaling cascades, including MAPK, which is known to be activated by BDNF (Fig. 5E). We then asked whether this enrichment was specific to ribosomal fraction, or whether it represented a transcriptional change (i.e. overrepresentation in the total mRNA). We then isolated the identity of mRNAs from two enriched GOs (“synapse organization” and “positive regulation of MAPK cascade”) and compared their log_2_FC with the log_2_FC from all mRNAs identified in the sequencing (Suppl. Fig. 4E-F). This comparison demonstrated that the enrichment is specific to the purified fraction, with no prominent changes found in the total mRNA (Fig. 5F-G). The GO analysis of mRNAs that were downregulated, on the other hand, revealed an enrichment of GOs related mostly to metabolic activities, such as “ribosome” or “mitochondrion organization” (Fig. 5H). When we compared the components from the GOs “ribosome” against the change of all mRNAs, we found a ribosomal fraction-specific depletion of ribosomal protein transcripts (Fig. 5I). Surprisingly, they were enriched in the total fraction, when compared against all mRNAs (Fig. 5J; Suppl. Fig. 4G). Comparison of the GO “microtubule” components did not show any significant enrichment in either fraction (Fig. 5J; Suppl. Fig. 4H).

To determine the identity of mRNAs that directly rely on GADD34 to be translated, we plotted the log_2_FC of all mRNAs shared in the sequencing of WT over KO cells (Fig. 5K). Again, overall distribution showed no obvious correlation between samples, but it allowed us to identify mRNAs that were upregulated (blue dots) or downregulated (orange dots) in WT cells exposed to BDNF, but not significantly changed in KO cells treated the same way (“corrected” mRNAs). GO analysis of the “corrected” mRNAs that were upregulated in the WT cells indicated that they were mostly related to either synaptic composition or synapse activity (Fig. 5L). The downregulated “corrected” mRNAs, on the other hand, were mostly related to mitochondrial activity or ribosomes (Fig 5M). Comparing the fold changes of the candidate GOs “synaptic organization” and “positive regulation of MAPK cascade” revealed that transcripts were enriched in the total fraction of KO cells versus WT, whereas the opposite was true for the purified fraction (Fig. 5N). On the other hand, comparison of the mRNAs that belong to the GO “ribosome” showed an enrichment in the total fraction of WT when compared to KO, and this relationship was inverted in the purified fraction. (Fig 5O). Again, the GO “microtubule” showed no detectable difference between fold changes of WT and KO (Fig 5O). Overall, these results reveal a central role for GADD34 in the BDNF-induced synthesis of proteins that are directly involved in synaptic plasticity and highlight that eIF2-dependent translation initiation is a novel pathway that mediates BDNF-dependent synaptic plasticity.

Importantly, BDNF stimulation have a significant effect in KO cells, as we found 2662 DEGs in the TRAP fraction (1162 upregulated, 1500 downregulated) and 411 in the total mRNA fraction (225 upregulated, 186 downregulated) (Suppl. Fig. 4I-J). Indeed, when we isolated IEGs fold changes, we found no obvious alteration in the induction of IEGs when compared to controls (Suppl. Fig. 4K; total mRNA fraction not shown). However, bioinformatic analysis revealed no outstanding GO being altered among upregulated mRNAs in the TRAP fraction. On the other hand, mRNAs related to the GO “post synapse” were downregulated after stimulation with BDNF. We then compared the fold changes between WT and KO cells of genes with the GOs “synaptic organization” (upregulated in WT) and “post synapse” (downregulated in KO) and found that the upregulation seen in the TRAP fraction was not present in the total mRNA fraction (Fig. 5P), suggesting dysfunction at the translational level. In summary, these results suggest that BNDF-induced plasticity relies on a translational program driven by the GADD34-eIF2α axis.

### BDNF increases interactions between G-actin and GADD34

The fact that GADD34 can rapidly promote eIF2α dephosphorylation following neuronal stimulation with BDNF indicates that this process could be integrated with early events involved in synaptic consolidation. Notably, to effectively bind to eIF2α, GADD34 must interact with G-actin, which stabilizes the GADD34-eIF2α interaction^89, 90^. Indeed, a SynGO analysis of mRNAs significantly changed in the TRAP-seq showed that the sub-group “actin cytoskeleton dynamics” is overrepresented (Fig 6A, highlighted quadrants). An in-depth analysis of the mRNAs showed no specific enrichment in the sequencing (Fig 6B). The change in *Cfl1*, however, caught our attention due to its central role in re-shaping the actin cytoskeleton during early synaptic plasticity^91^. We hypothesized that BDNF could promote actin destabilization through cofilin, leading to increased interactions between GADD34 and G-actin. Consistent with this idea, stimulation of neurons with BDNF for 1 hr resulted in a significant decrease in phosphorylated cofilin (Fig 6C-D). Surprisingly, though, this decrease is not due to changes in phosphorylation per se (Fig 6E), but rather due to increased amounts of total cofilin (Fig 6F).

**Figure 6.**
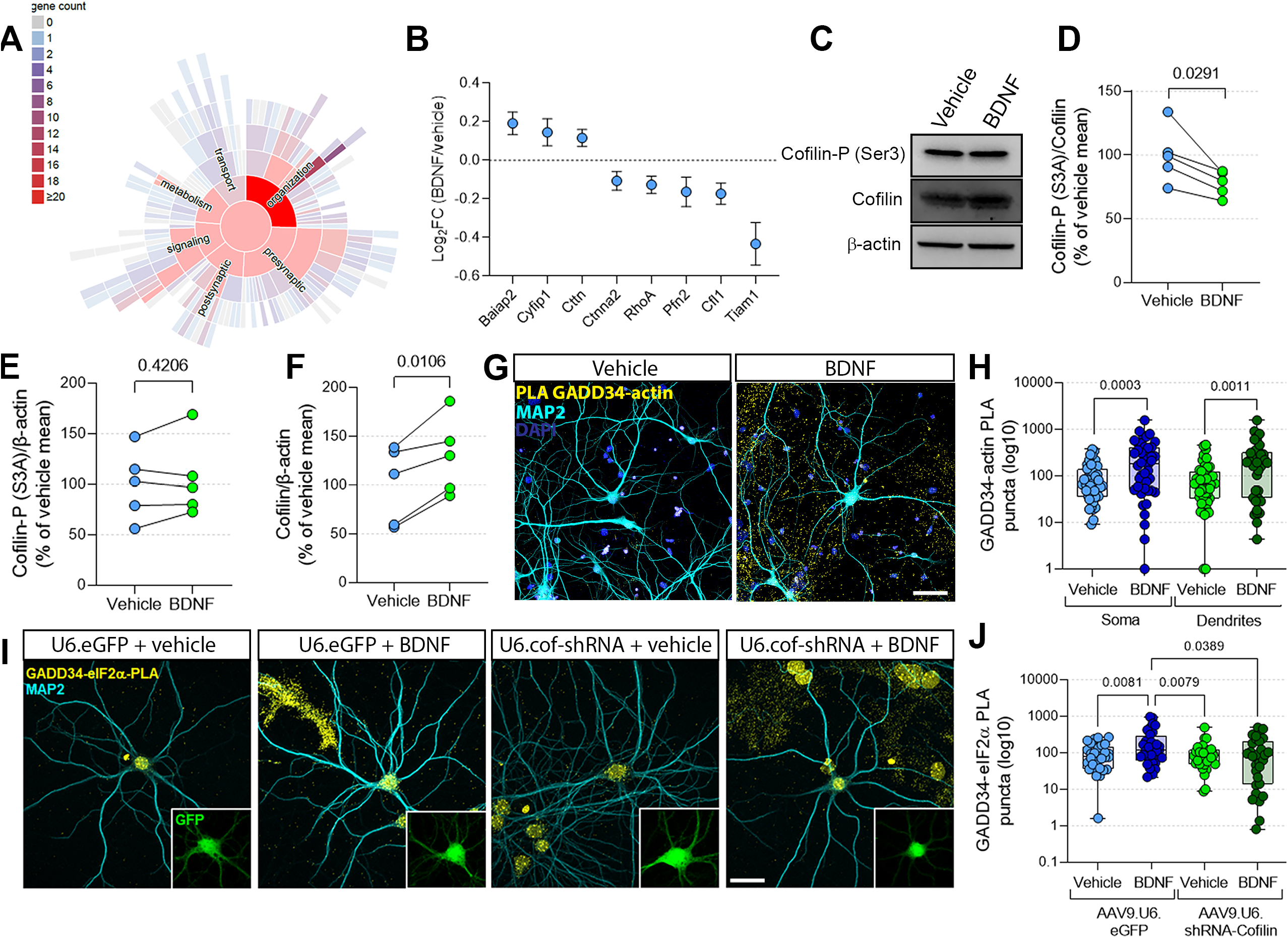
BDNF increases the physical interaction between G-actin and GADD34. (**A**) SynGO sunburst representative chart, highlighting the group “organization”, where the subgroup for cytoskeleton rearrangement is inserted (also highlighted). (**B**) Log_2_FC of changes of the mRNAs differentially loaded in ribosomes in response to BDNF that are related to the SynGO term analyzed. (**C**) Representative western blots probed cofilin-P (Ser 3, upper row), total cofilin (middle row) and β-actin (lower row) using samples from neurons exposed to vehicle or BDNF for 1 hr. (**D**) Quantification of the ratio of cofilin-P over cofilin (n=3 primary cultures). Paired *t* test. (**E**) Quantification of the ratio of cofilin-P over β-actin (n=3 primary cultures). Paired *t* test. (**F**) Quantification of the ratio of cofilin over β-actin (n=3 primary cultures). Paired *t* test. (**G**) Representative images of the PLA for GADD34-pan actin in primary neurons. Insets represent the PLA signal in the soma (black dots). Below the images are dendritic representations of the PLA signal. Yellow=PLA GADD34-eIF2α; Cyan=MAP2; Blue=DAPI. Scale bar=100 μm. (**H**) Quantification of panel **G** (n=39-44 neurons from 3 independent cultures). Statistical analysis was performed to compare differences in each compartment, independently. Unpaired *t* test. (**I**) Representative images of GADD34-eIF2α PLA in primary neurons transduced with AAV9.U6.shRNA-Cofilin or control, and then exposed to BDNF for 1h. Yellow=GADD34-eIF2α PLA; Cyan=MAP2. Scale bar=50 μm. (**J**) Quantification of panel **I** (n=29-31 neurons/group from 3 independent cultures). Two-Way ANOVA followed by Dunnet’s *post-hoc* test.

Because cofilin promotes the conversion of F-actin into G-actin, we asked whether BDNF could increase GADD34-actin interactions. A PLA showed that BDNF increased the interaction between both proteins (Fig 6G-H). To verify that the observed interaction was due to G-, rather than F-actin, we used latrunculin A to sequester G-actin, rendering it unavailable for protein interactions^92–94^. The sequestration of G-actin completely ablated the GADD34-actin PLA signal (Suppl. Fig. 5A-E). Finally, knockdown of cofilin with targeted shRNA (Suppl. Fig. 5C-D) blocked the increase in GADD34-eIF2α interactions induced by BDNF (Fig. 6I-J), without modifying total protein levels of GADD34 (Suppl. Fig. 5E-F). Overall, these results demonstrate that BDNF increases cofilin activity, facilitating physical interaction between eIF2α and GADD34.

### Increasing cofilin activity promotes eIF2α dephosphorylation and protein synthesis

Because BDNF decreased cofilin phosphorylation (Fig 6B-C) and promoted GADD34-G-actin interactions (Fig 6D-G) that mediates binding of GADD34 to eIF2α, we examined whether cofilin activity could act as a modulator of eIF2-dependent translation. To determine whether increased cofilin activity is sufficient to increase GADD34-eIF2α interactions, we transduced primary neurons with two viral vectors bearing overexpressing systems: AAV5.CamKIIα.cofilin-WT-HA (cof-WT) or AAV5.CamKIIα.cofilin-S3A-HA (cof-S3A) (Fig. 7A), where the former increases the amount of total cofilin inside the cell and the latter results in the overexpression of a hyperactive, phospho-dead mutant cofilin^95^ (Fig S6A-E). Using this system, we verified that the G/F-actin ratio was increased when cof-S3A was overexpressed (Fig S6F-G).

**Figure 7.**
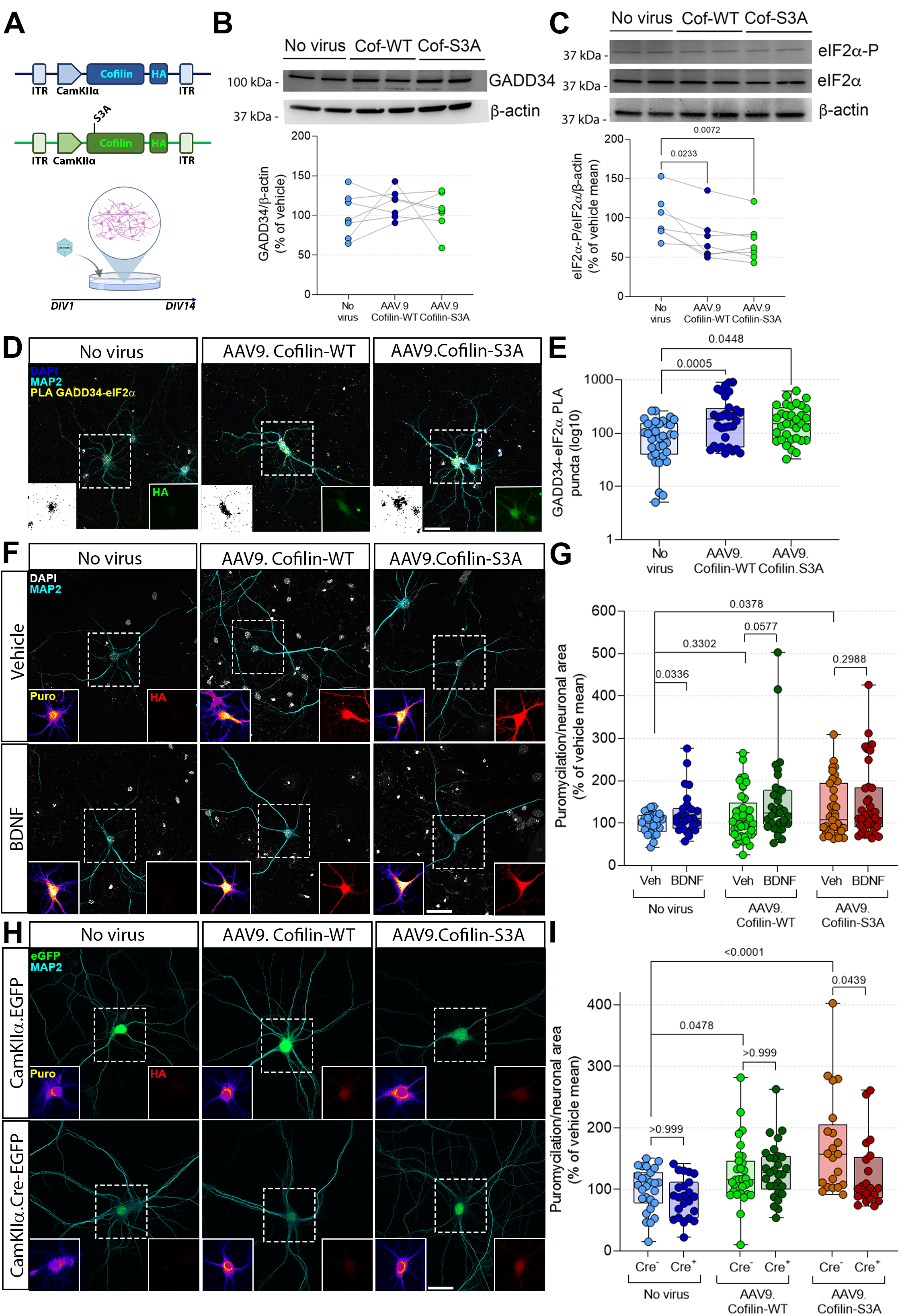
Increasing cofilin activity promotes protein synthesis and eIF2α dephosphorylation. (**A**) Viral system used to overexpress wild type or mutant cofilin in primary neurons. (**B**) Western blot probed for GADD34 (upper lane) or β-actin (lower lane) on samples from neurons transduced with either no virus, cof-WT or cof-S3A. Below is the quantification of GADD34 normalized by β-actin (n=5 independent primary cultures). One-way ANOVA followed by Dunnett *post-hoc* test. (**C**) Top panel: western blot probed for eIF2α-P (ser 5, upper lane), total eIF2α (middle lane) or β-actin (lower lane) on samples from neurons transduced with no virus, cof-WT or cof-S3A. Bottom panel: quantification of eIF2α-P normalized by eIF2α and β-actin (n=5 independent primary cultures). One-way ANOVA followed by Dunnett *post-hoc* test. (**D**) Representative images of GADD34-eIF2α PLA in neurons transduced with no virus, AAV5.CamKIIα.Cof-WT-HA or AAV5.CamKIIα.Cof-S3A-HA. Yellow=PLA; Cyan=MAP2; Blue=DAPI; Green=HA (lower right inset). Scale bar = 100 μm. (**E**) Quantification of the total PLA signal normalized by the neuronal area (n=32-34 neurons from 3 independent cultures). One-way ANOVA followed by Dunnett *post-hoc* test. (**F**) Representative images of *in situ* SUnSET in neurons transduced with no virus (left column), AAV5.CamKIIα.Cof-WT-HA (middle column) or AAV5.CamKIIα.Cof-S3A-HA (right column), exposed to vehicle (upper row) or BDNF (lower row). Cyan=MAP2; Gray=DAPI; Pseudo color=puromycin (lower left inset); Red=HA (lower right inset). Scale bar=100 μm. (**G**) Quantification of “E” (n=27-41 neurons from 3 independent cultures). Comparisons vehicle versus BDNF for each group (no virus, cof-WT or cof-S3A): unpaired *t* test. Comparisons between groups: One-way ANOVA followed by Dunnett’s *post-hoc* test. (**H**) Representative images of *in situ* SUnSET in neurons transduced with no virus (left column), AAV5.CamKIIα.Cof-WT-HA (middle column) or AAV5.CamKIIα.Cof-S3A-HA (right column), co-transduced with CamKIIα.eGFP (upper row) or CamKIIα.Cre-eGFP (lower row). Cyan=MAP2; Green=eGFP; Pseudo color=puromycin (lower left inset); Red=HA (lower right inset). Scale bar=50 μm. (**I**) Quantification of “H” (n=21-31 neurons from 3 independent cultures). Comparisons vehicle versus BDNF for each group (no virus, cof-WT or cof-S3A): unpaired *t* test. Comparisons between groups: Two-Way ANOVA followed by Dunnett’s *post-hoc* test.

We then determined whether the overexpression of either cof-WT or cof-S3A changed total levels of GADD34 and eIF2α-P. Expression of either type of cofilin did not alter expression of GADD34 (Fig. 7B), whereas both caused downregulation of eIF2α-P (Fig. 7C). These findings suggest that cofilin promotes increased GADD34-eIF2α interactions through an increase in available G-actin. We thus performed a PLA for GADD34-eIF2α in neurons transduced with either cof-WT or cof-S3A and found that overexpression of both forms of cofilin enhanced GADD34-eIF2α interactions (Fig. 7D-E). Our results led us to test whether cofilin overexpression could increase basal translation. Our results indicated that cof-S3A, but not cof-WT, significantly increased neuronal translation (Fig. 7F-G, light-colored points). However, when stimulated with BDNF, neurons transduced with cof-WT showed increased translation when compared with non-transduced neurons stimulated the same way (Fig. 7F-G, dark-colored points). Finally, we queried whether the cofilin-dependent increase in protein synthesis is mediated by GADD34. To investigate this, we obtained primary neurons from GADD34^fl/fl^ mice, and transduced them with two viruses: (1) CamKIIα.Cre, to ablate the expression of GADD34 in excitatory neurons; and (2) either no virus, cof-WT or cof-S3A. Strikingly, we found that the ablation of GADD34 blocked the increase in *de novo* protein synthesis mediated by cof-S3A (Fig. 7H-I). Altogether, these data demonstrate that increases the amount of available G-actin triggers a GADD34-dependent increase in protein synthesis.

## Discussion

Here, we interrogated the molecular mechanisms that contribute to fast changes in neuronal *de novo* translation due to activity. Among the conserved checkpoints for translation is the dephosphorylation of eIF2α, which is critical for many forms of long-lasting synaptic plasticity and memory consolidation^2^. Rapid dephosphorylation of eIF2α occurs in CamKIIα-expressing excitatory neurons and somatostatin-expressing interneurons in the hippocampus following contextual threat conditioning^78^ and in CamKIIα-expressing principal neurons in the lateral amygdala^77, 79^. Accordingly, rapid chemogenetic-induced and transient increases in eIF2α-P during early stages of memory consolidation can impair that process^79^. Our results indicate that translation of GADD34 and cytoskeleton dynamics converge to promote activity-dependent eIF2α dephosphorylation within an hour, thereby driving increases in neuronal translation, including the synthesis of proteins closely related to synaptic remodeling and consolidation.

### Plasticity-related dephosphorylation of eIF2α relies on GADD34 activity

Despite demonstration that dephosphorylation of eIF2α contributes to long-lasting plasticity and long-term memory formation, a molecular mechanism for eIF2α dephosphorylation in these contexts had not been identified. Herein, we provide evidence that GADD34, a scaffolding protein that mediates eIF2α dephosphorylation by PP1^96^, is responsible for rapid BDNF-induced dephosphorylation of eIF2α. Our findings offer an attractive explanation as to how BDNF shapes the neuronal proteome shortly after activation. We show that BDNF increases the levels of GADD34 and decreases eIF2α-P levels in primary neurons. Furthermore, BDNF promotes the physical interaction of GADD34 with eIF2α, (Fig 4), which is required for decreased levels of eIF2α-P. We cannot exclude the possibility that simultaneous events are occuring upstream to silence the eIF2α kinases given that PKR, GCN2, and PERK (all eIF2α kinases) have been shown to mediate learning and memory^97–102^. We propose that GADD34 rapidly promotes eIF2α dephosphorylation in neurons, but the downregulation of eIF2α kinases represent a long-term solution to keep translation at a higher rate during late phases of consolidation. This is supported by the notion that GADD34 has a high turnover rate due to the Δpest domains in its structure^96^, although these are thought to modulate PP1 activity, rather than promoting GADD34 degradation^103^. Moreover, we cannot also rule out an orchestrated act between GADD34 and CReP to drive translation upregulation during memory consolidation, as both proteins were shown to have complementary effects in the unfolded protein response^104^, and CReP has been demonstrated to have a role in synaptic plasticity and long-term memory formation^105^. Further studies are necessary to determine whether neuronal eIF2α dephosphorylation following activity is, in fact, a multi-factorial process, and whether it is a long-lasting or acute episode triggered by neuronal activation.

Understanding the molecular pathways that underlie activity-dependent translation initiation and their subcellular localization is key to unravel the initial steps of neuronal response to pre-synaptic stimulation. Proteomic studies suggest that BDNF increases the amount of translation initiation factors in dendritic compartments^106^. In addition, previous studies showed that a rapid, but transient, MNK1-mediated phosphorylation of eIF4E occurs when synaptodendrosomes were exposed to BDNF, which was correlated with an upregulation of *de novo* translation^49, 65^. Remarkably, our TRAP-seq revealed that effectors and targets of the ERK and mTORC1 pathways seem to be consistently modulated by BDNF through GADD34, as demonstrated by the changes in translating mRNAs related to translation and MAPK signaling, indicating a cross-regulation between these translation-promoting pathways. Notably, there is no evidence suggesting that eIF2 subunits are enriched in dendrites after potentiation, which suggests a different method of translation regulation^64, 67, 106^. The dephosphorylation of eIF2α triggered by a rapid, local increase in GADD34 levels could explain this and deems closer examination.

In addition to the modulation of translation factors, mRNA translation is heavily impacted by *trans* components, such as RNA-binding proteins (RBPs). RBPs bind to *cis-*regulatory elements present in both coding and non-coding sequences and give stability to mRNAs^16, 107^, allowing them to travel long distances within the cell without risking degradation by cellular machinery, and mediate active mRNA transportation to sub compartments^108^. However, RBPs also repress translation through different mechanisms, including inhibition of the preinitiation complex (CPEBs, FMRP and CYFIP1)^109, 110^, the 80S complex formation (ZBP1)^111^ or even translation elongation^112, 113^. Notably, RBPs are an integral part of mRNP granules that are distributed along dendrites and are sensitive to BDNF stimulation^53, 114–116^. It is unclear whether eIF2α dephosphorylation triggers the translation of mRNP-contained mRNAs. Our TRAP-sequencing data, thus, warrants deeper investigation to interrogate RBPs that are differentially modulated by BDNF in a translation initiation-dependent manner.

### Translational regulation of GADD34 expression in neurons

Historically, GADD34 expression has been associated to the ISR^73, 75^. In contrast, GADD34 expression increases in neuroblastoma cells when stimulated for 6 hr with a cannabinoid receptor agonist, which is reliant on CREB-dependent gene transcription^82^. Our findings are similar in that they demonstrate a stress-independent increase in GADD34 protein levels. However, our findings differ from the previous studies in neuroblastoma cells in that we found no differences in GADD34 mRNA levels. Instead, we found that the BDNF-induced increase in GADD34 levels is regulated at the translational level. A previous study found that *de novo* translation of GADD34 can be achieved through chemical LTP induction in acute hippocampal slices^117^. Nevertheless, it should be noted that there are differences in the temporal window for measuring GADD34, extracellular stimuli delivered, and the cell types utilized in the two studies, which could explain the mechanistic differences for how GADD34 levels are increased.

As mentioned above, most previous studies have examined GADD34 expression during conditions of cellular stress when its expression is mediated by ATF4 at transcriptional level^73, 118, 119^. However, the GADD34 mRNA contains two uORFs, components of the 5’ UTR of mRNAs that were originally described to negatively impact ribosome scanning of the coding sequence^120, 121^. In fact, the second uORF presents a Pro-Pro-Gly motif juxtaposed to the stop codon, which is poorly translated and can thus induce ribosome stalling upstream to the coding sequence^122–125^. Substituting this motif by Ala-Ala-Ala releases the brake in GADD34 translation, and elevated expression can be found even under resting conditions^120^. Therefore, it is possible that BDNF triggers the release of this brake of ribosomes and translation re-initiation in neurons, which would explain the increase in GADD34 expression we observed in our experiments. In fact, ribosome foot printing analysis of neuronal transcripts revealed that uORF-containing mRNAs are actively translated in both soma and dendrites^24^, building on the challenging of the notion that uORFs only suppress translation^74^. In fact, *PPP1R15A* mRNA contain two short uORFs, which are associated with higher possibility of reinitiation of translation at the main ORF^126–128^. Additionally, we cannot exclude the possibility that *PPP1R15A* mRNAs are stored in mRNA granules, membrane-less structures present in dendrites and axons that contain quiescent ribosomes and mRNAs. Granule components can be rearranged after neuronal stimulation and trigger translation of these transcripts locally, ensuring a rapid response to activity^129^. The fact that we can identify *PPP1R15A* mRNA in dendrites, and that BDNF increases its association to polysomes support this hypothesis. Nevertheless, more experiments are required to understand how exactly uORFs modulate *PPP1R15A* mRNA translation, and what type of control takes place in neurons.

### GADD34 expression converges with cytoskeleton dynamics to promote eIF2α dephosphorylation

We found that BDNF can promote an increase in the interaction between GADD34 and actin, which acts as a stabilizer of the primary interaction between GADD34 and eIF2α. Our experiments suggest that this interaction relies on an increased availability of G-actin after neuronal stimulation, through decreased phosphorylated cofilin after exposure to BDNF. The role of cofilin during synaptic consolidation is controversial. Classically, LTP is associated with higher phosphorylation of cofilin and elevated levels of actin polymerization, a mechanism that appears to rely on protein synthesis^130, 131^. On the other hand, the induction of LTD drives the opposite effect, with less phosphorylated cofilin and higher actin depolymerization^132^. This dichotomy was challenged by another report that described a massive translocation of active cofilin to the dendritic spine at early stages of LTP^133^. Previous reports suggested that cofilin is transiently dephosphorylated by the phosphatase slingshot 1 at early stages of synapse activation to sever F-actin and generate a source of G-actin^134–136^. Moreover, actin monomers are crucial for insertion of AMPA-Rs at the surface of dendritic spines after LTP. In fact, dephosphorylated cofilin stimulates the integration of AMPA-Rs to the surface, wheras the phosphorylated version has the opposite effect^137^. Because eIF2α-P can influence the amounts of AMPA-R present in the postsynaptic density^29^, and because we found that artificially increasing G-actin levels can promote GADD34-eIF2α interactions, it is possible that actin monomers increase AMPA-Rs at the synapse by promoting eIF2α dephosphorylation.

Our findings indicate that the overexpression of cofilin can promote neuronal protein synthesis, providing a link between the dynamic regulation of the cytoskeleton with an increase in translation. The interplay between actin dynamics and protein synthesis is critical for neuronal function^34–36, 138^, and when unbalanced appears to underlie neuronal dysfunction in fragile X syndrome (FXS). In FXS model mice, the lack of fragile X protein results in increased Rac1 activity, a GTPase that promotes actin remodeling, and increases eIF4E-eIF4G interactions, thus potentiating *de novo* translation^110^. Here, we show an additional molecular pathway through which actin dynamics can regulate neuronal translation initiation by promoting the interaction between GADD34-eIF2α. A next step of investigation requires more detailed analysis of structural changes in actin in the dendritic spine and the consequences for local translation initiation at this subcellular structure.

In conclusion, our findings demonstrate a novel molecular pathway through which neurons can increase protein synthesis upon BDNF stimulation. This pathway relies on the release of a translational brake on GADD34, which in turn can bind to eIF2α-P in both soma and dendrites in a G-actin-dependent manner to promote *de novo* protein synthesis (Suppl Fig. 7). These events culminate with the translation of mRNAs that encode proteins involved in synaptic plasticity. Our results open a new avenue towards understanding the molecular mechanisms of translational control that underlie long-lasting changes in neuronal function that are induced by neuronal activity that contribute to memory formation and that are altered in cognitive disorders.

## Methods

### Reagents

For the full list of reagents used, catalog numbers, and distributors, please refer to Table 1.

### Primary neuron preparation and maintenance

E17 pregnant mice were obtained and euthanized by cervical dislocation. The embryos were removed from the vitellin sack and brains were extracted and kept on ice-cold HBSS + 0.37% glucose. Cortico-hippocampal masses were isolated from the brain stem and cerebellum with use of blunted-end tweezers, and meninges were carefully removed, avoiding tissue puncturing. Tissue was kept on ice during the whole process. For chemical digestion, tissue was incubated for 5 min with 0.25% trypsin solution at 37°C. Trypsin was inactivated with two sequential washes with DMEM + 10% FBS and 1% Pen/Strep. For mechanical isolation of cells, tissue was resuspended in supplemented DMEM and disrupted with fire-polished Pasteur glass pipettes (15 strokes). Cells then were centrifuged at 900 *g*/5 min/RT and carefully resuspended in Neurobasal supplemented with 2% B27, 1% Pen/Strep and 1% GlutaMax at 37°C. The total amount of cells in suspension was determined by manual count using a Neubauer chamber. For imaging analysis, cells were plated at a final density of 100,000 cells/ well (24-well plates and 4-well culture slides) or 200,000 cells/well (12-well plates). For western blotting and polysome profiling analysis, cells were plated at a density of 2,000,000 cells/well (6-well plates) or 20,000,000 cells/well (Petri dishes). Cells were plated in glass coverslips or polypropylene plastic plates coated with a 0.1% poly-L-ornithine solution supplemented with 4 µg/ml of laminin. After 1 hr, the Neurobasal was aspirated with vacuum from the wells and replaced with fresh Neurobasal, supplemented as above. Cells were always kept at 37°C/5% CO2 conditions, and 40% of media was replaced every three days with fresh Neurobasal, until *DIV* (days *in vitro*) 14, when cells were used for experimentation. For experiments with either BDNF or thapsigargin, cells were treated at the final day of culture with a final concentration of 50 ng/ml of BDNF or 0.5 µg/ml of thapsigargin (times of incubation are delineated in the figure legends). For puromycilation and puro-PLA experiments, cells were exposed to 0.5 µg/ml of puromycin 5 min prior to fixation.

### RNA scope

Cells were cultured in chambered slides and treated/fixed as described above. To perform fluorescent *in situ* hybridization, the protocol of the manufacturer was followed. Briefly, cells were permeabilized for 5 min using protease IV diluted 1:15, rinsed 3x in PBS. Hybridization was performed for 2 hr at 40°C using ACD Bio HyBEZ oven. Cells were washed 2x with 1x RNA scope wash buffer, and then amplification was performed with the following conditions: AMP 1-FL for 30 min at 40°C, AMP 2-FL for 15 min at 40°C, AMP 3-FL for 30 min at 40°C and AMP 4-FL for 30 min at 40°C. Between each amplification step, cells were washed twice with 1x RNA scope wash buffer. After RNA scope, immunostaining was performed. Cells were blocked with 5% normal goat serum solution and incubated for 30 min at room temperature with anti-MAP2 antibody. Cells were washed 2x with 1x RNA scope buffer and incubated with secondary anti-chicken for 30 min at room temperature. Cells were washed again 2x with RNA scope buffer, rinsed in Mili-q water and mounted using Antifade Prolong mounting media + DAPI.

### Immunocytochemistry

After treatment, Neurobasal was aspired with a vacuum and briefly rinsed with pre-warmed PBS. PBS was aspired and cells were fixed for 20 min with PBS + 4% paraformaldehyde (PFA) + 4% sucrose solution at RT. Fixative solution was removed and cells were rinsed 4x with PBS. For permeabilization, neurons were incubated for 15 min/RT with PBS + 0.5% Triton X-100 and rinsed twice with PBS. The coverslips then were incubated for 1 hr with blocking solution consisting of 5% NGS in PBS and then incubated overnight/4°C with primary antibodies diluted in blocking solution. The next day, cells were washed 3×5 min with PBS and incubated for 1 hr at room temperature with secondary antibodies diluted in blocking solution, then washed 3x 5 min with PBS again. Coverslips were mounted on Prolong + DAPI, when 405 channel was available for DAPI staining. Otherwise, Prolong without DAPI was used instead. Slides were incubated for 24 hr at room temperature to dry mounting media before imaging. The concentrations of the primary and secondary antibodies are shown in Table 1. Imaging was done using a Leica SP8 Confocal microscope within a week after finishing immunocytochemistry. A 63X magnification objective was used, and 10-15 z-stacks were obtained per image, separated by 0.5 µm. Images were obtained at a final resolution of 512 x 512 pixels.

### Western blotting

Cells were scraped from the plate in cold RIPA buffer (10 mM Tris-Cl pH8.0, 1 mM EDTA, 1% Triton X-100, 0.1% sodium deoxycolate, 0.1% SDS, 140 mM NaCl) and flash frozen on dry ice. The resulting lysates were sonicated and cleared through centrifugation (10.000 *g/*10 min/4°C). Supernatant was transferred to a new tube and total protein mass was quantified using BCA kit. Samples were prepared in Laemmli buffer supplemented with 10 mM DTT and boiled at 95°C for 3 min. After cooling to room temperature, samples were spun and applied in 4-20% Tris-Glycine electrophoresis gels to a final mass of 20-40 µg of protein per lane, depending on the experiment performed. Proteins were separated at a fixed voltage of 100V, and then transferred to polyvinylidene difluoride (PVDF) membranes using iBlot2 apparatus (23V/6 min). Membranes then were incubated with 5% milk for 30 min, then 5% BSA for 30 min. Membranes were incubated with primary antibodies diluted in 5% BSA at 4°C/overnight/rocking, and then washed 3x with TBS + 0.1% Tween-20 (TBS-T). Membranes were incubated with secondary antibodies diluted in 5% BSA for 1hr at room temperature while rocking, and then washed 3x with TBS-T and 3x with TBS. Development was done with use of ECL kits from Amersham/GE, using a Protein Simple apparatus (Kodak). For concentrations of primary and secondary antibodies used, please refer to table 1.

### Polysome profiling

Cultured mouse cortical neurons at DIV 14 were treated with either vehicle or BDNF (50 ng/ml) for 1 hr. Neurons then were treated with cycloheximide (CHX) at 100 µg/ml for 1 min at 37°C. After washing twice with phosphate buffered saline with 100 µg/ml CHX-treated cells we scraped in polysome buffer (20 mM Tris pH 7.5, 150 mM NaCl, 5 mM MgCl_2_, 100 μg/ml cycloheximide, 1 mM DTT) and centrifuged at 300 *g*/5 min/4°C in a swing bucket rotor. Pellets were lysed using polysome lysis buffer (20 mM Tris pH 7.5, 150 mM NaCl, 5 mM MgCl_2_, 1% Triton-X 100, 24 U/ml Turbo DNase, 100 U/ml RNasin Plus, 100 μg/ml cycloheximide, 1 mM DTT, protease inhibitor cocktail, and 8% glycerol) and the lysate was triturated with 23-gauge syringe, followed by centrifugation 16.100 *g*/10 min/4°C. Samples then were loaded onto 14 ml 10 to 50% sucrose gradients prepared in polysome buffer (20 mM Tris pH 7.5, 150 mM NaCl, 5 mM MgCl2, 100 μg/ml cycloheximide, 1 mM DTT, 100U/ml RNasin Plus, and 8% glycerol) and centrifuged for 2 hours 45 min at 36,000 revolutions per minute (rpm) at 4°C in a SW41 Ti swing-out rotor. Polysome profiling was carried out using a Piston Gradient Fractionator (Biocomp) with continuous monitoring at 254 nm using a Model EM-1 Econo UV detector (BioRad). Monosome and polysome fractions of approximately 400 μl were collected with a fraction collector (Gilson FC203B).

### RNA purification and RT-PCR

TRAP and total lysate RNA were isolated via the RNeasy Plus Kit (Qiagen). The samples were first lysed and homogenized in Buffer RLT Plus. The lysates then were centrifuged through a gDNA Eliminator spin column for 30 sec. The flow through was saved and 300 µL of 70% ethanol was added. The sample then was passed through the RNeasy spin column for 15 seconds. RNA bound to the membrane and the contaminants were washed away. The RNA in the spin column membrane was washed in 700 µL of buffer RW1 and 1000 µL of buffer RPE. The isolated RNA was eluted in 30 µL of RNase free water. The concentration of RNA for each sample was determined using NanoDrop and was later normalized to 10 ng/µL. The RNA expression levels of the target mRNA *PPP1R15A* along with an endogenous control gene, *GAPDH*, were measured in a qPCR reaction. The qPCR reactions were performed using the Luna Universal One-Step RT-qPCR kit. For each reaction, 19 µL of a qPCR master mix containing Luna Universal One-Step Reaction mix, Luna WarmStart RT Enzyme mix, Forward Primer, Reverse Primer, and Nuclease-free water was added to a 96 well plate. 1 µL of the 10 ng RNA sample was then added to the 96 well plate, and 1 µL of RNase free water was used as the negative control. Technical replicate qPCR reactions were run on the same plate. Next, the qPCR reactions were run using SYBR fluorescence on a BioRad CFX96 qPCR cycler. Following the qPCR reactions, the threshold values (Ct) were used to calculate the fold change. First, the Ct value of the endogenous control was subtracted from the Ct value of the target gene to determine the target ΔCt. (*Target* Δ*Ct = Ct_GADD34_ - Ct_GAPDH_*). The average ΔCts from all three of the technical replicates was then calculated along with the average ΔCts from the negative control. The negative control average ΔCt values then were subtracted from the target ΔCt values giving a ΔΔCt value (ΔΔCt = ΔCt*_Target_*-ΔCt*_NTC_*). Finally, to determine the fold change, 2 was raised to the negative ΔΔCt (Fold Changes = 2^-ΔΔCt^). For the polysome profiling fraction qPCRs, RNA collected from polysome profiling experiments was isolated from the sucrose gradient via the Direct-zol RNA MiniPrep Kits (Zymo). The RNAs from each fraction were lysed and homogenized in 400 µL TRI-Reagent (Zymo). An equal volume of 95-100% ethanol was added to the lysed samples and thoroughly mixed. The mixture then was transferred to a Zymo-spin IICR Column and centrifuged. The RNA bound to the spin column membrane and the flow through was discarded. The spin column was washed with 800 µL of RNA Pre-wash and 700 µL of RNA wash buffer. The RNA then was eluted in 30 µL of nuclease-free water. The concentration for each RNA fraction was determined using Nanodrop and later normalized to 10 ng/µL. RT-qPCR reactions were performed as described previously.

### Proximity Ligand Assays

Cells were rinsed, fixed, and blocked as sdescribed above. Proximity Ligand Assays (PLA) were performed following instructions from the manufacturer. Briefly, cells were incubated with either a combination of anti-GADD34 (rabbit)/anti-eIF2α (mouse), anti-GADD34 (rabbit)/anti-actin (mouse), anti-GADD34 (rabbit)/anti-puromycin (mouse) or anti-eIF2αP (rabbit) overnight/4°C. In all experiments, cell also incubated with anti-MAP2 (chicken) and, when transduced with viruses, anti-mCherry (rat). Cells were washed 3×5 min with PBS and incubated at 37°C for 1 hr with PLA probes (rabbit or mouse) diluted 1:5 in blocking buffer, and secondary anti-Chk (1:500). Cells were washed 4×5 min/RT with 1x PLA buffer A. Cells then were incubated at 37°C for 30 min with ligase diluted in 1x PLA ligase buffer, and subsequently washed 4x 5 min at room temperature with 1x PLA buffer A. Finally, for amplification, cells were incubated at 37°C for 100 min with polymerase diluted in 1x PLA amplification buffer and rinsed twice with buffer B at room temperature. Cells then were washed 2x 10 min at room temperature with buffer B, 3x 5 min with PBS and mounted with Prolong + DAPI. After the mounting media was dry, cells were immediately imaged using a Leica SP8 Confocal microscope. A 40X magnification objective was used, and stacks were obtained separated by 0.2 µm (25-40 stacks per image). Images were obtained at a final resolution of 1024 x 1024 pixels.

### Translating Ribosome Affinity Purification (TRAP)

TRAP was performed as previously described (Heiman *et al.*, 2014). On the day before sample collection, the affinity matrix (composed of protein L-coated magnetic beads + anti-GFP antibodies) was prepared. Briefly, MyOne T1 Dynabeads were incubated for 30 min with 120 µg of protein L at RT, rotating. Beads were washed 5x with RNAse-free PBS + 3% protease and IgG-free BSA and incubated with 50 µg of each anti-GFP antibody (19C8 and 19F7) for 1h/RT/rotating. Beads were washed 3x with low-salt buffer (20 mM HEPES KOH pH 7.3, 150 mM KCl, 10 mM MgCl_2_, 1% NP-40, 0.5 mM DTT, 100 µg/ml Cycloheximide and protease/phosphatase inhibitor) and stored until use. On the next day, samples from neurons treated for 1 hr with either vehicle or 50 ng/ml BDNF were collected. Prior to sample collection, neurons were exposed to 100 µg/ml cycloheximide for 1 min to block translation. Cells were washed 3x with ice-cold RNAse-free PBS + 100 µg/ml cycloheximide on ice, and gently scraped in cell lysis buffer (100 mM HEPES KOH pH 7.3, 750 mM KCl, 50 mM MgCl_2_, 1% NP-40, 0.5 mM DTT, 40 units/ml RNAsin, 20 units/ml Superasin, 100 µg/ml cycloheximide and protease/phosphatase inhibitor). The lysates were transferred to a pre-chilled glass homogenizer and 12 strokes were performed with an automatic pestle, avoiding sample aeration. Cell lysates were transferred to pre-chilled 1.5 ml RNAse-free tubes and centrifuged at 2000 *g*/10min/4°C. The supernatant was transferred to a new tube and supplemented with 1/9^th^ of the volume of 300 mM DHPC solution (e.g. 111 µl for every ml of sample). Samples were homogenized by inversion and incubated for 5 min on ice, followed by new centrifugation at 20,000 *g*/10min/4°C. Supernatant was collected and incubated at 200 µl of affinity matrix/sample overnight/4°C/rotating. On the next day, beads were isolated using a magnet, and washed 4x with high-salt buffer (20 mM HEPES KOH pH 7.3, 350 mM KCl, 10 mM MgCl_2_, 1% NP-40, 0.5 mM DTT, 100 µg/ml cycloheximide and protease/phosphatase inhibitor). After the last wash, buffer was removed, and beads were allowed to return to room temperature before adding 300 µl of lysis buffer from RNEasy Miniprep Plus kit supplemented with β-mercaptoethanol. Samples were incubated for 10 min at room temperature to allow elution, and beads were removed magnetically, and supernatant transferred to a new tube.

### RNA sequencing

TRAP was performed as described above. Quality control of both total RNA and purified RNA was measured using Agilent HS TapeStation. cDNA then was synthesized with Takara SMART-Seq HT kit with 1 ng of RNA input that had RIN of 5 and greater via Agilent HS TapeStation. Libraries were constructed with 0.25 ng of cDNA input via Nextera XT DNA library kit. cDNA and libraries amplification had both 12 cycles. Quality control of libraries were analyzed via Agilent HS TapeStation and Invitrogen Quant-it. Samples were pooled at equal molarity before sequencing. Final libraries were sequenced paired-end 50 cycles on an Illumina NovaSeq6000 S1 100 cycle Flow Cell-v1.5 with 1% Phix spike-in.

### Bioinformatics

For alignment of Fastq files, sequencing adapters and low-quality bases were trimmed using Cutadapt version 1.8.2 using the following arguments (-q 20 -O 1 -a CTGTCTCTTATA)^139^. Trimmed reads were then aligned to the mouse genome (mm10) using STAR version 2.7.7a^140^ and raw gene counts were obtained using the argument (*-quantmode*).

After alignment, a count matrix was generated using scripts generated in house. The count matrix was batch normalized using ComBat-Seq^141^, and used as input for DESeq2 package for differential expression analysis^142^. In the analysis design, both genotype (WT vs KO) and treatment (vehicle vs BDNF) were included. Data was transformed using variance stabilizing transformations (vst) and used as input for PCA. For the PCA, the plotPCA function (imbued in the DESeq package) was used. For unsupervised clustering, the DESeq object generated was transformed into a matrix, which then was used to calculate a correlational matrix between samples. Heatmaps were generated using pHeatmap package. Log_2_ fold changes were retrieved using lfcShrink function, using “apelgm” as an estimator^143^. To obtain a list of significant DEGs, the list of genes was subset for FDR < 0.1, as previously described^144^.

Gene Ontology analysis was performed using clusterProfiler package^145^. Briefly, the list of significantly upregulated or downregulated DEGs were transformed into ENSEMBL IDs using bitr function, and the list was queried against a background list, that consisted of all genes identified on the sequencing. All GO types were analyzed together, with a *p* value cutoff of 0.05. The *p* value was adjusted using Benjamin & Hochberg’s method. GOs included in figures were selected based on (1) significance; and (2) groups of interest. synGO analysis^146^ was performed using the official repository (www.syngoportal.org) using standard methods.

### G/F-actin ratio experiments

G/F-actin fractionation was performed as previously described^110^. After cells were treated with either vehicle or BDNF, cell lysates were scraped in ice-cold “G-actin buffer” (10 mM KHPO_4_, 50 mM KCl, 2 mM MgCl_2_, 1 mM EGTA, 0.2 mM DTT, 0.5% Triton X-100, 1 mM sucrose and 100 mM NaF, supplemented with protease/phosphatase inhibitors). Lysates were kept ice-cold through the whole process. Cell lysates were sonicated and spun at 15.000 *g*/30 min/4°C. Supernatant, containing G-actin, was saved and frozen until use. The pellet was resuspended in ice-cold F-actin buffer (1.5 mM guanidine hydrochloride, 1 mM sodium acetate, 1 mM CaCl_2_, 1 mM ATP and 20 mM Tris-HCl pH 7.4, supplemented with protease/phosphatase inhibitors) and sonicated again. Lysates were centrifugated at 15,000 *g*/30 min/4°C and supernatant containing F-actin was saved. Protein samples were quantified and loaded on tris-glycine gels, as described above. A western blot using anti-pan-actin as the primary antibody was performed to develop the total levels of actin in each sample.

### Image analysis

All fluorescence intensity, PLA, RNA scope and western blot band density analysis were performed using Fiji (Image J). For fluorescence intensity and RNA scope, the channel containing MAP2 (a neuronal marker), GFP or mCherry (when neurons were transduced with viral constructs) was initially isolated and used as template for a mask to define the region of interest (i.e. neuronal area). This mask was applied over the channel containing the staining of interest, and a threshold was applied to separate real signal from noise, using staining performed with no primary antibodies as background. The integrated density then was obtained and normalized by the total neuronal area. For PLA, an ImageJ-based script developed by Heumuller and colleagues^33^ was kindly provided by Dr. Erin Schuman and used to calculate total intensity of punctate signal. For western blots, the bands were straightened using rotation tool and then total staining density was calculated using the raw integrated density parameter in Fiji. For every image analysis, the experimenter was blind to all conditions. No parameters of brightness or contrast were altered prior to quantification.

### Data analysis

Data analysis and plotting were performed using GraphPad Prism (v9). Details regarding the tests used for each experiment are described in figure legends. Paired analysis were used for Western blot analysis. Unpaired analysis were used to analyze immunofluorescence, RNA Scope and PLAs. When three independent conditions were analyzed together, One-Way ANOVA was performed. When two interacting conditions (i.e. genotype vs treatment) were analyzed, Two-Way ANOVA was performed. For time course data, One-Way ANOVA repeated measures was performed. All graphics display minimum to maximum value, with median represented as a black line in the barplot. Data normality was visualized using Q-Q plots before appropriate statistical test application. Outliers were excluded from the analysis using the Grubb’s test (α = 0.05) after all experimental data was collected.

## Author contribution

M.M.O. and E.K. conceptualized and designed the project. M.M.O., M.K.E., M.M., K.B.M., M.M., E.H.L., E.A.G., N.N. and S.C. designed, performed, and analyzed experiments. M.M.O. and E.K. wrote and edited the manuscript. T.A. and E.K. contributed animals, materials, and analysis tools.

## Figure legends

**Supplementary Figure 1. Presence of *PPP1R15A* mRNA in neurons.**

(**A**) RNA scope-based fluorescent in situ hybridization (FISH) targeting GADD34 mRNA (punctate signal) in primary neurons exposed to either vehicle or 1 μM thapsigargin for 1h. Green = FISH; Blue = DAPI; Cyan = MAP2. Scale bar = 50 μm.

(**B**) Representation of the radial quantification of *PPP1R15A* mRNA expression in primary neurons.

(**C**) Radial quantification of GADD34 mRNA puncta in neurons, from soma (light blue background) to dendritic portions (dark blue background) (n=3 independent primary cultures).

(**D**) Total amounts of *PPP1R15A* mRNA in either soma or dendrites of neurons exposed to either vehicle (blue bars) or (red bars) 1 μM thapsigargin for 1 hr (n=33-34 neurons/condition, collected from three independent primary cultures). Statistical analysis was performed to compare differences in each compartment, independently. Unpaired *t* test, * = p<0.05; **** = p<0.0001. Mean ± SEM.

(**E**) Total percentage of cells transduced with AAV9.CamKIIα.Cre, AAV9.EF1a.mCherry.SICO-shRNA.PPP1R15A, or both.

(**F**) Percentage of cells successfully co-transduced with AAV9.CamKIIα.Cre and AAV9.EF1α.mCherry.SICO-shRNA.PPP1R15A.

**Supplementary Figure 2. Validation of methods to study translation of *PPP1R15A* mRNA in neurons.**

(**A**) Low-magnification representative images of primary neurons co-transduced with AAV9.CamKIIα.Cre and AAV5.CAG.FLEX.L10a-eGFP viruses. Cyan = MAP2; Blue = DAPI; Green = eGFP. Scale bar = 200 μm.

(**B**) Quantification of panel A (n = 5-6 tilescans per experiment, 2 independent primary cultures), depicting the percentage of primary neurons successfully co-transduced with AAV9.CamKIIα.Cre and AAV5.CAG.FLEX.L10a-eGFP viruses.

(**C**) Timeline of treatments with cycloheximide (10 μg/ml) and puromycin (0.5 μg/ml) to validate the puro-PLA approach in neurons.

(**D**) Representative images of puro-PLA targeting *de novo* translation of GADD34. Cyan = MAP2; Blue = DAPI; Yellow = Puro-PLA. Scale bar = 50 μm.

(**E**) Quantification of “E” (n = 9-20 neurons, obtained from 3 independent experiments). One-way ANOVA followed by Dunnet *post-hoc* test, *** = p<0.001; **** = p<0.0001. Mean ± SEM.

**Supplementary Figure 3. Validation of GADD34-eIF2a PLA and *de novo* protein synthesis measurement using puromycin.**

(**A**) Representative images of GADD34-eIF2α PLA performed in primary neurons exposed to either vehicle or 1 μM thapsigargin (1h or 3h). Cyan = MAP2; Blue = DAPI; Yellow = GADD34-eIF2α PLA. Scale bar = 50 μm.

(**B**) Quantification of “A” (n = 8-16 neurons, obtained from 2 independent experiments). One-way ANOVA followed by Dunnet *post-hoc* test, ns = not significant; ** = p<0.01. Mean ± SEM.

(**C**) Representative images of neurons exposed to cycloheximide (10 μg/ml) and puromycin (0.5 μg/ml) to validate the puromycilation approach as a measure of *de novo* translation in neurons. Scale bar = 100 μm.

**Supplementary Figure 4. Validation of TRAP-Seq.**

(**A**) Experimental design to purify ribosomes of paired WT and KO cells.

(**B**) Principal component analysis of the samples obtained from TRAP-Seq.

(**C**) Unsupervised clustering of the samples obtained from TRAP-Seq.

(**D**) Volcano plot highlighting the cell type-specific markers in the purified samples.

(**E-H**) Volcano plots highlighting the components of the GOs “synaptic organization”, “MAPK signaling”, “Ribosomes” and “Microtubules”.

(**I - J**) Volcano plot evidencing DEGs between KO and WT cells, in total lysate (I) or purified samples (J). Orange = upregulated. Blue = downregulated.

(**K**) Fold changes of IEGs comparing WT and KO samples.

**Supplementary Figure 5. Characterization of GADD34-actin PLA and AAV9.U6.Cofilin-shRNA.**

(**A**) Representative images of the PLA for GADD34-actin in primary neurons exposed to 200 nM of latrunculin A for 5 hr, followed by exposure to 50 ng/ml of BDNF for 1 hr. Green = PLA GADD34-actin; Red = MAP2; Blue = DAPI. Scale bar = 50 μm.

(**B**) Quantification of “F” (n=32-37 neurons, collected from three independent primary cultures). Two-Way ANOVA followed by Tukey’s *post-hoc* test, **** = p<0.0001. Mean ± SEM.

(**C**) Representative western blots of samples transduced with either AAV9.U6.GFP or AAV9.U6.Cofilin-shRNA. Upper row = Cofilin; lower row = β-actin.

(**D**) Quantification of **C** (n = 3 independent primary cultures). One-Way ANOVA followed by Dunnet *post-hoc* test. Ns = non-significant; *** = *p*<0.001. Mean ± SEM.

(**E**) Representative western blots of samples transduced with either AAV9.U6.GFP or AAV9.U6.Cofilin-shRNA. Upper row = GADD34; lower row = β-actin.

(**F**) Quantification of **E** (n = 3 independent primary cultures). One-Way ANOVA followed by Dunnet *post-hoc* test. Ns = non-significant. Mean ± SEM.

(**G**) Representative western blots of samples transduced with either AAV9.U6.GFP or AAV9.U6.Cofilin-shRNA. Upper row = eIF2α-P; middle row = eIF2α; lower row = β-actin.

(**H**) Quantification of **G** (n = 3 independent primary cultures). One-Way ANOVA followed by Dunnet *post-hoc* test. Ns = non-significant. Mean ± SEM.

**Supplementary Figure 6. Characterization of cofilin overexpression systems.**

(**A**) Representative western blots of samples transduced with either no virus, AAV5.CamKIIα.Cof-WT-HA (middle column) or AAV5.CamKIIa.Cof-S3A-HA (right column). Upper row = Cofilin-P (Ser3). Middle row = Total cofilin. Lower row = HA tag.

(**B**) Quantification of total cofilin-P levels, normalized by β-actin (n=3 independent primary cultures). One-way ANOVA, One-way ANOVA followed by Dunnet *post-hoc* test, ns = not significant; * = p<0.05. Mean ± SEM.

(**C**) Quantification of total cofilin levels, normalized by β-actin (n=3 independent primary cultures). One-way ANOVA, One-way ANOVA followed by Dunnet *post-hoc* test, ns = not significant; * = p<0.05. Mean ± SEM.

(**D**) Quantification of cofilin-P/cofilin (n=3 independent primary cultures). One-way ANOVA, One-way ANOVA followed by Dunnet *post-hoc* test, ns = not significant; * = p<0.05. Mean ± SEM.

(**E**) Percentage of cofilin-P originated from endogenous vs viral expression.

(**F**) Percentage of cofilin originated from endogenous vs viral expression.

(**G**) Representative western blots of G/F-actin fractionation assays. Upper row = G-actin; Middle row = F-actin; Lower row = HA tag.

(**H**) Quantification of “G” (n=6 independent primary cultures). One-way ANOVA followed by Dunnet *post-hoc* test, ns = not significant; * = p<0.05. Mean ± SEM.

