## Supplementary figures and images for "The Integrated Stress Response effector GADD34 is repurposed by neurons to promote stimulus-induced translation"

### Supplemental Figures

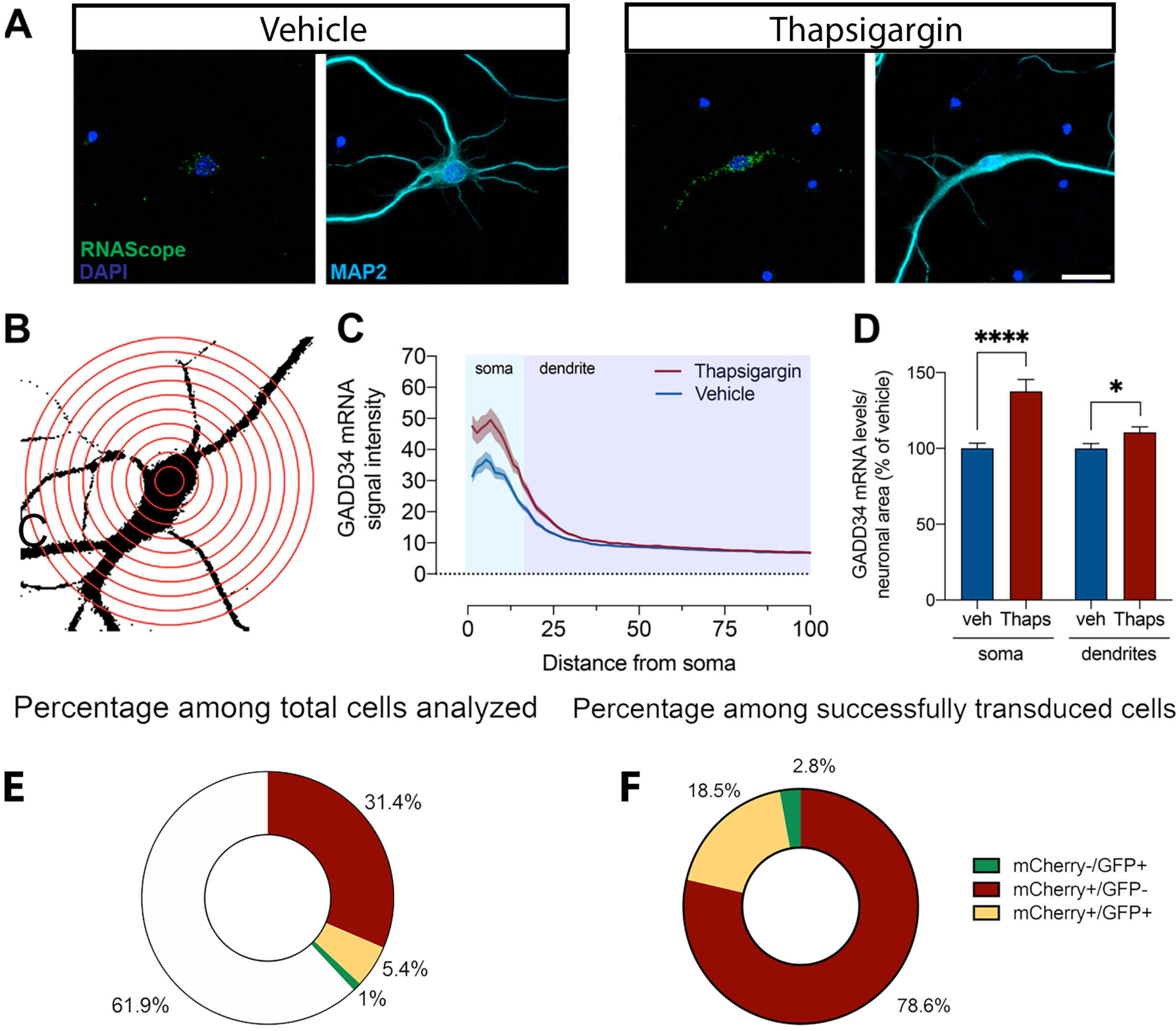

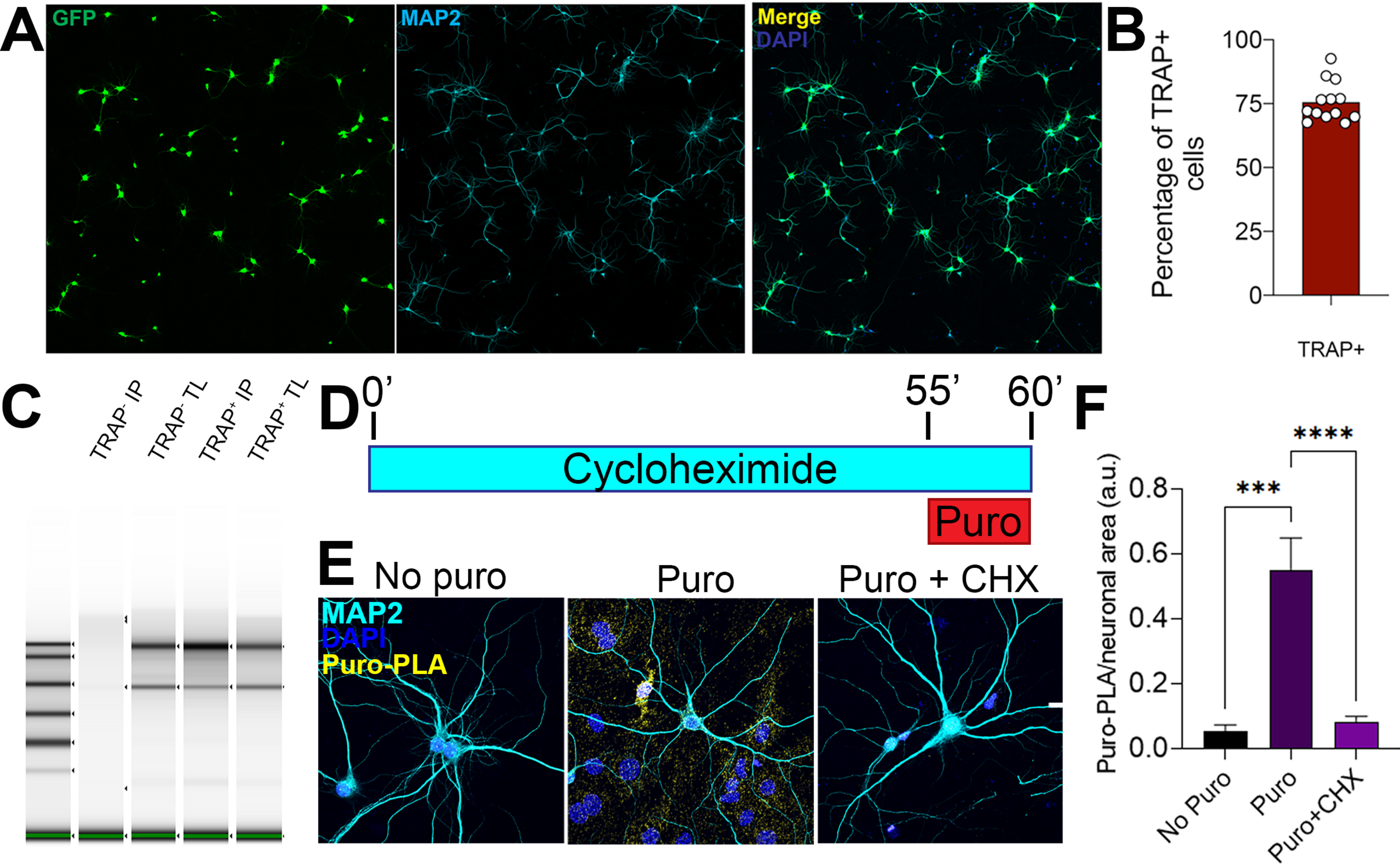

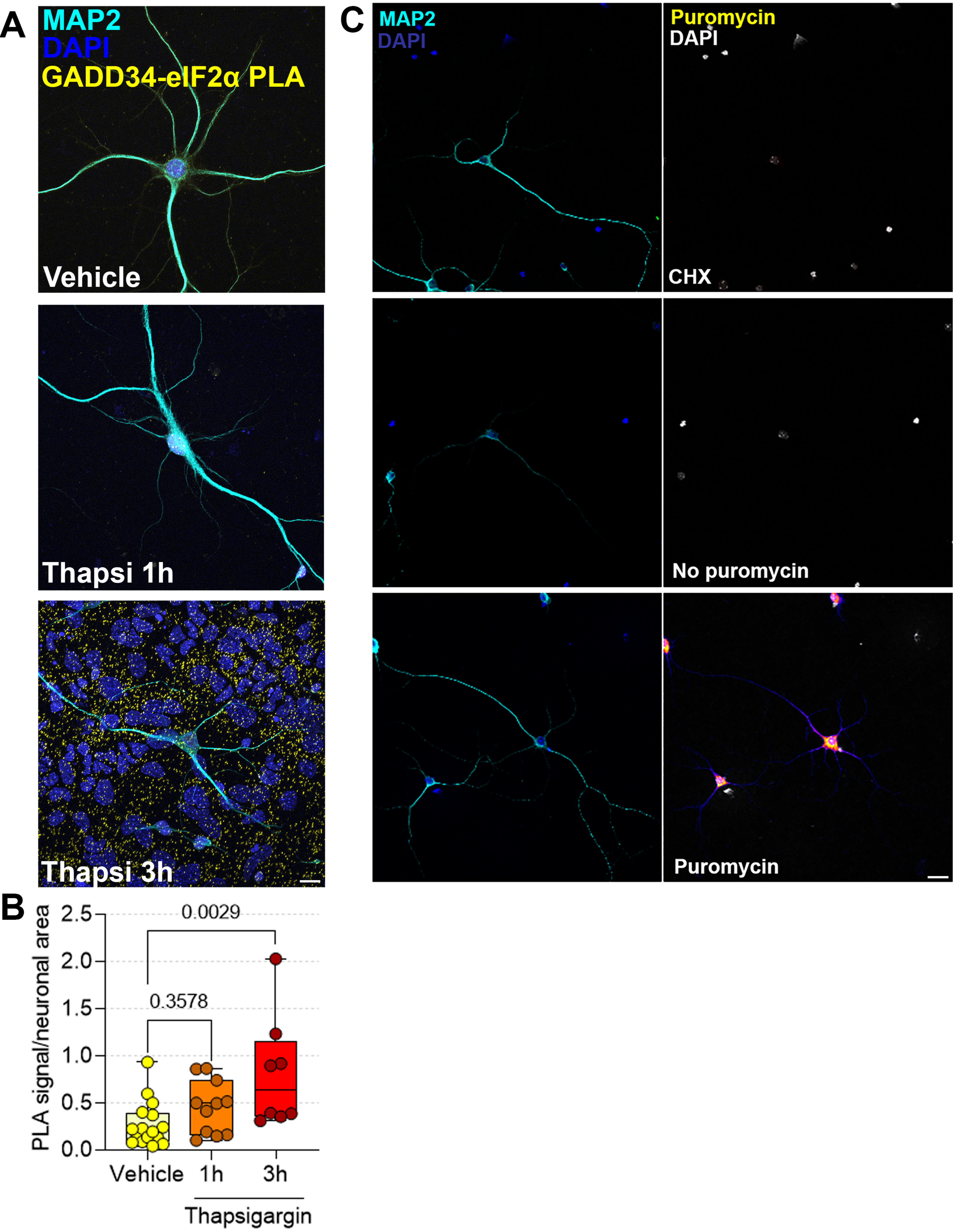

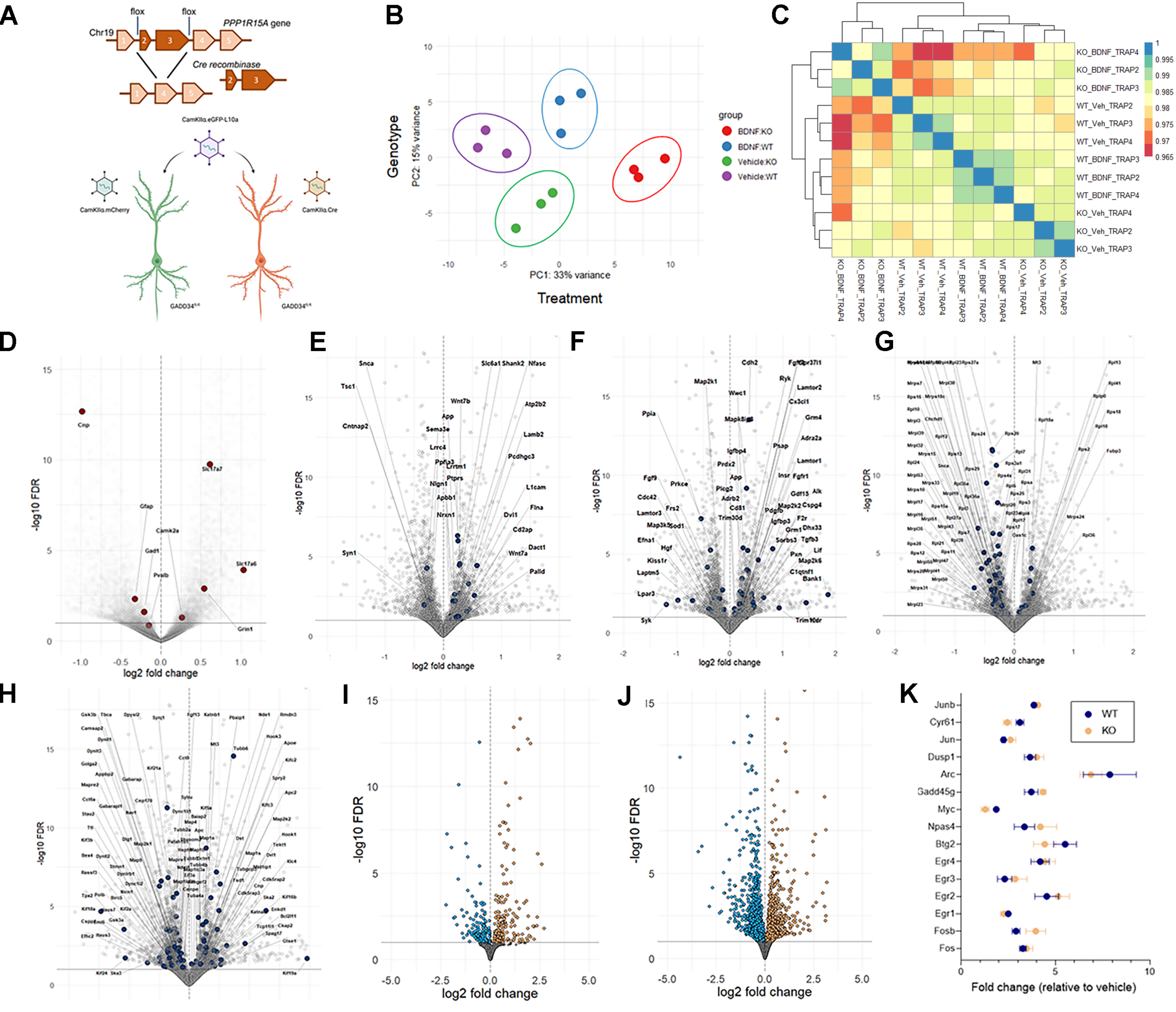

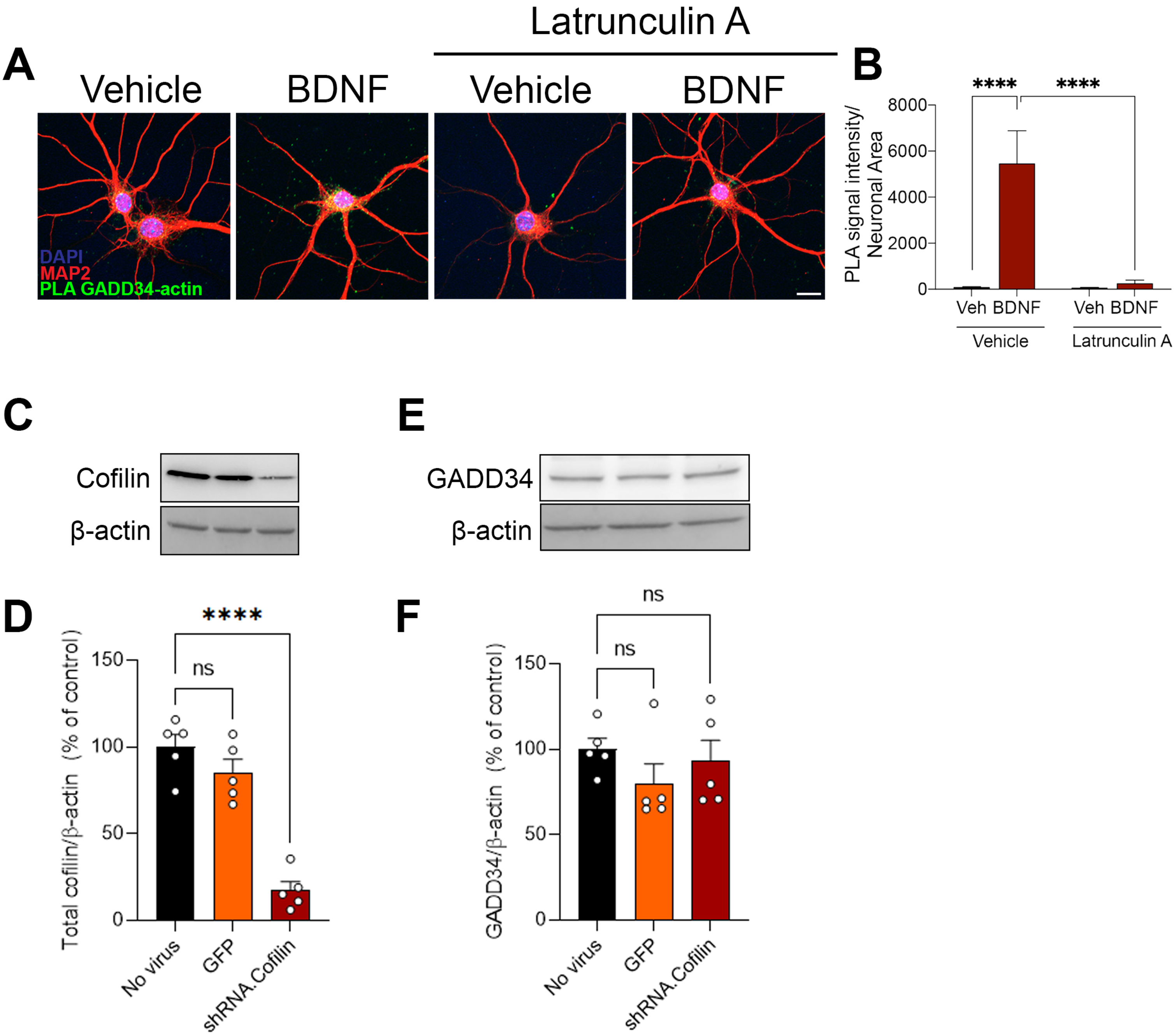

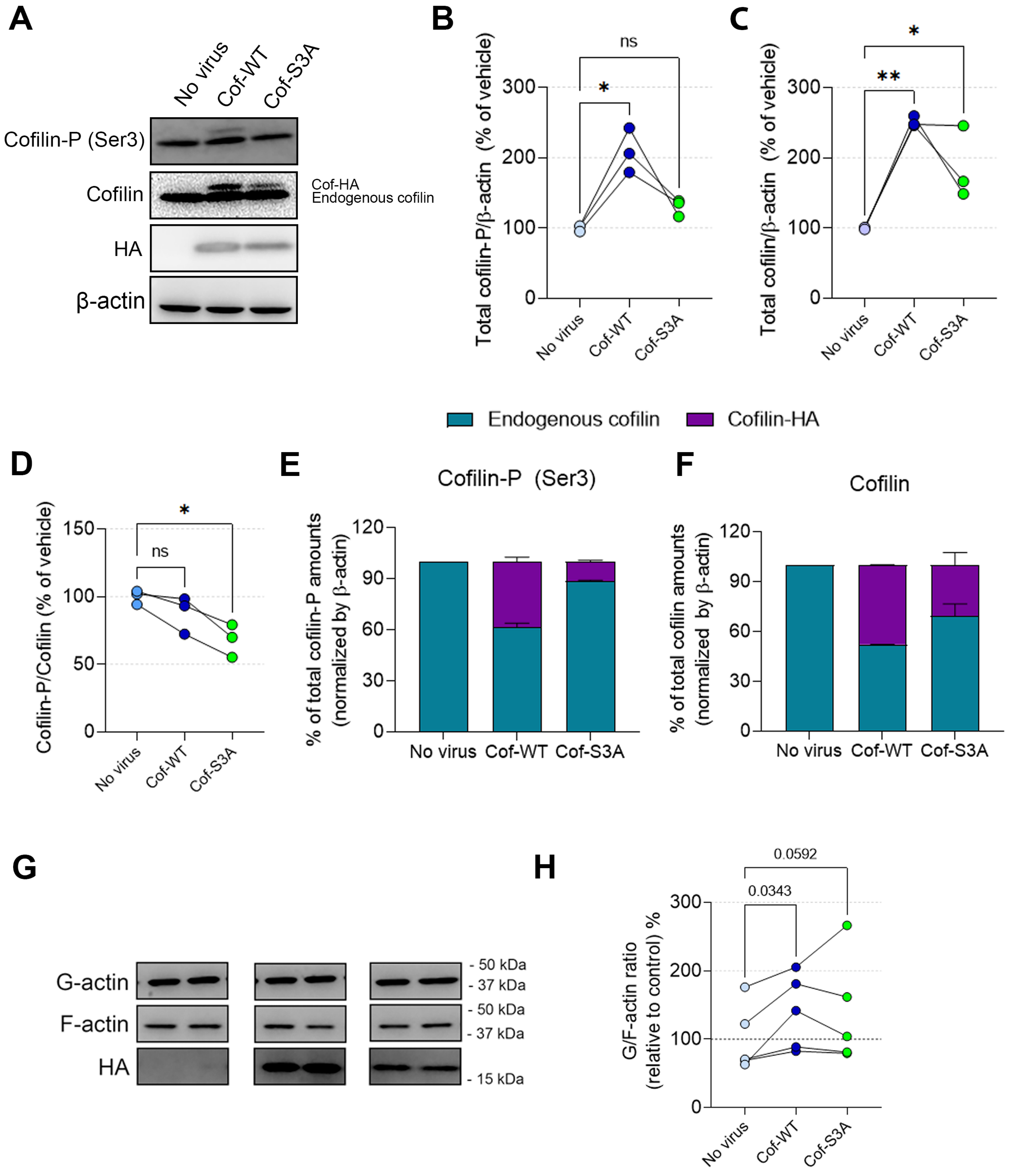
